# Nuclear encoded photosynthesis genes are specifically controlled by the NuA4 complex

**DOI:** 10.1101/846592

**Authors:** Tomasz Bieluszewski, Weronika Sura, Anna Bieluszewska, Michał Kabza, Mateusz Abram, Piotr Włodzimierz, Wojciech Dzięgielewski, Maja Szymańska-Lejman, Catherine Lachance, Nancy De Winne, Geert De Jaeger, Jacques Côté, Jan Sadowski, Piotr A. Ziolkowski

## Abstract

NuA4, an essential histone acetyltransferase complex, is required for efficient transcription in eukaryotes. Using genome editing, genomic approaches and biochemical assays, we characterized plant homologues of two key components of this complex, EPL1 and EAF1 in *Arabidopsis thaliana*. Surprisingly, we found that loss of AtEPL1, which is necessary for enzymatic activity of NuA4, is not lethal. Contrary to yeast, mutants lacking AtEAF1, responsible for complex targeting, display severe pleiotropic phenotype which copies that of *Atepl1*. *Atepl1* and *Ateaf1* mutants grow slowly, contain reduced chlorophyll levels and small chloroplasts. We provide evidence that these alterations are not caused by disturbance of GLK transcription factors, the major regulators of chloroplast development. Using ChIP-seq we show that H4 acetylation levels are dramatically reduced in the chromatin of the *Atepl1* mutant, while H3 acetylation remains mostly unchanged. We use our data to define NuA4-dependent genes and show that chloroplast-related genes are significantly overrepresented in this group, consistent with the pale-green phenotypes of the mutants. We propose that NuA4 was adopted in plants to control nuclear-encoded photosynthesis genes.

**Significance:** Photosynthesis depends on chloroplast proteins, most of which are nucleus-encoded and thus subject to control mechanisms common across eukaryotes. Here we show that NuA4, an evolutionary conserved transcriptional coactivator, is necessary for proper development of photosynthetic apparatus. Surprisingly, in contrast to yeast and metazoans, plants engineered to lack core NuA4 subunits are capable of vegetative development despite dramatic genome-wide loss of NuA4-dependent H4K5 acetylation. This chromatin perturbation seems to directly affect 350 genes which, in addition to reduced H4K5ac levels, display decreased transcript levels but no loss of transcription-related H3K9ac. A significant proportion of these genes are related to chloroplast function, particularly to translation, an intriguing parallel to the yeast NuA4’s role in transcription of ribosome biogenesis-related genes.

## Introduction

Substantial biochemical and genetic evidence support a functional interplay between chromatin structure and transcription in eukaryotes. The reverse genetic approach, possible due to advances made in yeast and animal models, provided a framework to our understanding of this relationship in plants. It is now evident that evolutionarily conserved enzymes and chromatin marks have been adopted in plants for functions in development, stress response and genome integrity. In fact, conserved chromatin transactions play central roles in regulation of plant-specific processes such as transition to flowering(1, 2) and flower morphogenesis(3, 4).

One of the characteristic features of plants are chloroplasts which carry out photosynthesis(5). As most of the chloroplast proteins are encoded by nuclear genes, chromatin should in principle have an influence on their expression. Molecular and genetic studies suggest that such a link may indeed exist in plants. Some observations showed that changes in the acetylation status of histones H3 and H4 over genes encoding plastidic proteins coincide with changes in light and nutrient availability as well as developmental stage(6, 7). Likewise, increased levels of H3K9 acetylation (H3K9ac) accompany increased transcriptional activity of these genes(8). In addition, mutations which disrupt genes involved in histone modifications and chromatin remodeling such as *AtGCN5*, *HAF2* (one of the two *Arabidopsis thaliana* genes encoding TAF1 homologues) or *BRM*, affect transcript levels of genes encoding plastidic proteins(9, 10). Interestingly, chromatin perturbations described so far produce mild photosynthetic or photomorphogenic phenotypes, despite often extreme impact on other processes(11). Here we show that the expression of the photosynthetic apparatus of a model plant *A. thaliana* depends on histone H4 acetylation by a highly conserved transcriptional coactivator, the Nucleosomal Acetyltransferase of H4 (NuA4) complex.

The budding yeast NuA4 complex consists of 13 different subunits which fall into 3 functional categories: enzyme, histone posttranslational modification reader and scaffold(12). Scaffold proteins Enhancer of Polycomb-Like 1 (EPL1), Esa1-Associated Factor 1 (EAF1) and Transcription-Associated 1 (TRA1) recruit reader subunits and transcription factors, greatly improving specificity of the acetyltransferase subunit Essential SAS2-related Acetyltransferase 1 (ESA1) towards histones H4, H2A and H2A.Z in the chromatin context(13–16). Molecular studies in yeast and animals support critical roles for the NuA4-dependent histone acetylation in DNA-repair and transcription(17, 18). Plant homologues of NuA4 subunits were linked by genetic studies to gametogenesis, ABA-responses, control of flowering time, chlorophyll synthesis, cell growth and ploidy(19–22). Some of these functions of the plant NuA4 have been confirmed to be mediated by acetylation of H4(21, 23, 24).

In this work, we introduce a set of genetic tools which gave us a unique opportunity to explore the consequences of complete loss of NuA4 integrity without compromising viability as in the case of the catalytic subunit mutants described in eukaryotic models before(19, 25, 26). Mutants lacking homologues of the scaffold subunits EPL1 or EAF1 consistently display pleiotropic phenotype, much more severe than the NuA4-mutant phenotypes described in plants so far. We show that the photosynthetic component of this phenotype is similar in magnitude, but different in origin, to the effects observed in the absence of the Golden2-Like (GLK) transcription factors which specifically promote chloroplast development in various plant species(27). Our genetic and transcriptomic analyses argue against NuA4 and GLKs acting in the same pathway. Instead, we show that loss of NuA4 integrity leads to dramatic global decrease of H4K5 acetylation with almost no impact on H3K9ac as well as widespread disruption of nuclear transcription. Our results suggest a direct link between NuA4 activity and expression of the GLK-independent nuclear genes encoding plastid ribosomal proteins(28) (npRPGs).

## Results

### NuA4 supports chloroplast development

The budding yeast NuA4 complex consists of two modules, the catalytic module called piccolo NuA4 (later picNuA4) and the regulatory module attached to picNuA4 through the platform subunit EAF1(12) (Fig. 1A). The EPL1 subunit plays a double role in NuA4 by enabling picNuA4 formation and binding to the regulatory part of the complex(15). Whereas the former function of EPL1 is essential, the latter is not(14). Consistently, the yeast strain lacking the C-terminal Enhancer of Polycomb B (EPcB) domain of EPL1 through which it binds to EAF1, phenocopies strains lacking functional EAF1(15). EAF1 and EPL1 are not only essential for the integrity of the NuA4 complex but also specific to NuA4, unlike most other subunits which are shared with other protein complexes, therefore, they are key to understanding NuA4 function(15, 29). Despite growing interest in the plant NuA4 complex, little is known about plant homologues of these two proteins.

**Figure 1.**
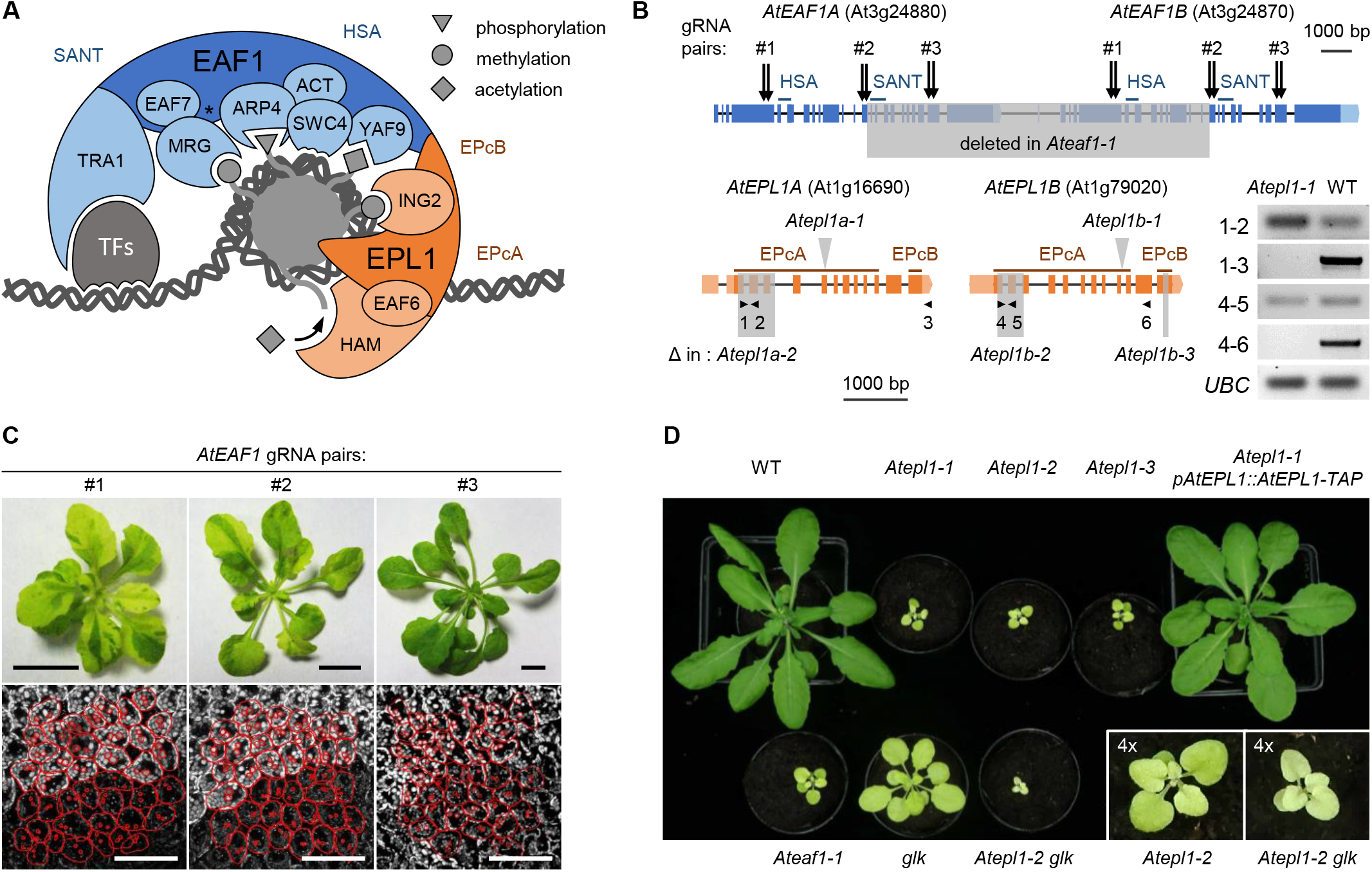
*Ateaf1* and *Atepl1* mutants as tools for genetic dissection of the plant NuA4 complex. A. Schematic overview of the NuA4 complex engaging chromatin. The catalytic module picNuA4 and the regulatory module are colored orange and blue, respectively. The asterisk between EAF7 and ARP4 indicates the lack of an EAF5 homologue in plants. Approximate positions of the crucial protein domains are indicated by the labels outside the shapes representing the subunits. B. Gene structure diagrams of the *AtEAF1* and *AtEPL1* loci. Target sites for the three double-gRNA constructs (gRNA pairs) used for *AtEAF1* mutagenesis are depicted by arrows. Shaded areas delimit Cas9-gRNA-induced deletions (Δ) characterizing particular alleles. Grey arrowheads indicate T-DNA insertion sites. Small black arrowheads represent primers used for the RT-PCR presented in the panel on the right. C. Transgenic plants carrying the three different Cas9-gRNA constructs. The scale bar length is 10 mm in all images. Lower panels are confocal images of chlorophyll autofluorescence in the palisade mesophyll cell layer of a leaf, centered on the border regions between green (upper part) and pale green (lower part) sectors. Red outlines delimit selected cells and chloroplasts. The scale bar is 50 μm in all micrographs. D. 4 week-old WT and mutant plants grown in the same conditions. The insets show 4x magnifications of the same image.

Our previous study of Arabidopsis EAF1 (AtEAF1) could not reveal its full importance due to the lack of a double mutant(24). Here, in order to generate a loss-of-function mutant, we disrupted both *AtEAF1* genes simultaneously by Cas9-gRNA (Fig. 1B). During screening for plants carrying heritable mutations, we observed the effects of sectorial somatic double knock-out of *AtEAF1* which was manifested as green/pale green mosaicism (Fig. 1C). Microscopic observation of the border regions between green and pale green sectors revealed a reduction in the size of chloroplasts in the pale green (presumably EAF1-negative) sectors (Fig. 1C), suggesting that AtEAF1 function is cell-autonomous. We observed the Cas9-gRNA induced mosaicism in plants carrying all 3 independent Cas9-gRNA combinations although the construct targeted downstream of the Swi3, Ada2, N-Cor and TFIIIB (SANT) domain produced a weaker effect, suggesting a minor role of the C-terminal part of AtEAF1 in chloroplast development (Fig. 1C). The association between the chloroplastic phenotype and the loss of AtEAF1 was confirmed in the *Ateaf1-1* homozygous mutant which was evenly pale green (Fig. 1D). In this mutant line, the 11-kb deletion produced a chimeric *AtEAF1A/B* gene containing a frameshift upstream of the SANT domain (Fig. 1B, Supplementary Fig. 1). The SANT domain is vital to the function of yeast EAF1(15), therefore, we consider *Ateaf1-1* a loss-of-function mutant and a reference for reverse genetics of the plant NuA4 complex.

*A. thaliana* contains two EPL1 orthologues which have been copurified with other plant NuA4 subunits previously(24, 30) but have not been studied genetically. In order to investigate the function of AtEPL1, we generated a double T-DNA insertion mutant (later *Atepl1-1*) by crossing two publicly available lines. Insertions in At1g16690 (*Atepl1a-1*) and At1g79020 (*Atepl1b-1*) are located within EPcA (*Atepl1a-1*, Fig 1B) or between EPcA and EPcB domains (*Atepl1b-1*, Fig. 1B). Whereas full-length transcripts are undetectable in the double homozygous mutant, both loci still produce aberrant transcripts encoding a polypeptide homologous to the N-terminal part of EPL1 (Fig. 1B), which is sufficient for picNuA4 activity in budding yeast but not for the binding to the regulatory part of the complex(15) (Fig. 1A).

Since the yeast EPL1 serves two functions in the NuA4 complex, one of which is potentially preserved in *Atepl1-1*, we used Cas9-gRNA to generate additional alleles of the *AtEPL1* genes. In order to obtain null alleles, we introduced frame-shifting deletions in the 5’ parts of the coding regions of both genes (Fig. 1B, Supplementary Fig. 2 and 3). As lethal effect of a double homozygous mutation was expected, we first generated two single homozygous mutants (*Atepl1a-2* and *Atepl1b-2*) and crossed them to obtain the *Atepl1-2* double mutant. In addition, we generated an in-frame deletion spanning the EPcB domain of AtEPL1B (*Atepl1b-3*, Fig. 1B, Supplementary Fig. 3) and crossed the mutant to *Atepl1-2* to obtain the *Atepl1a-2 Atepl1b-3* mutant (later referred to as *Atepl1-3*). Since the genomic deletion present in the *AtEPL1A* locus in the *Atepl1-2* mutant starts in an exon and ends in an intron, the exact transcript sequence cannot be predicted with confidence. In order to check if splicing of the *Atepl1a-2* allele leads to a frameshift, we sequenced 6 independent cDNA clones derived from RNA isolated from a homozygous *Atepl1-2* mutant. All clones displayed identical splicing pattern of the truncated intron IV which led to a frameshift, therefore this allele is unlikely to produce even partially functional AtEPL1A protein in significant amounts (Supplementary Fig. 2). Importantly, the strong pleiotropic *Ateaf1*-like phenotype shared by the *Atepl1* homozygotes was fully rescued in *Atepl1-1* homozygotes expressing tagged AtEPL1B (pAtEPL1B∷AtEPL1B-TAP, Fig. 1D), confirming loss of AtEPL1 as the cause of the phenotypic effects observed in *Atepl1* mutants.

### AtEPL1 is not essential for viability but key to NuA4 function

Contrary to our predictions based on homology to yeast, where EPL1 is an essential gene, all double *Atepl1* mutants described above were viable in double homozygote form, albeit clearly more defective than other plant NuA4 subunit mutants characterized to date (Supplementary Fig. 4). The viability of the *Atepl1* mutants gave us a unique opportunity to compare the relative contributions made by the EAF1-dependent NuA4 holocomplex and EPL1-dependent picNuA4 in plants. We used four quantitative phenotyping assays to describe and compare the growth- and photosynthesis-related phenotypes of *Ateaf1* and *Ateapl1* mutants. Additionally, we included in our analyses the well-characterized *glk1-1 glk2-1* mutant (referred to as *glk* below) in which cell-autonomous reduction in chloroplast size as well as significant reduction in rosette diameter have been observed previously(27, 31).

As an approximation for chloroplast- and cell-size we used surface projection areas of the organelles or cells obtained through live confocal imaging of the palisade mesophyll cell layer of rosette leaves. In order to eliminate variability resulting from changes in chloroplast size during leaf growth, we measured chloroplasts in fully expanded 4^th^ leaves of 10-true-leaf-rosettes. We found both chloroplasts and cells to be significantly smaller in all mutants compared to wild type (WT) plants (Fig. 2A, Supplementary Table 1A). The NuA4 mutants were indeed similar to the *glk* mutant in terms of chloroplast size but less so in terms of cell size (Fig. 2A). Notably, the *Atepl1-2* null-mutant displayed stronger reduction in chloroplast size than the *Ateaf1-1* mutant, possibly owing to residual non-targeted activity of picNuA4 in this line. It remains unclear why *Atepl1-3* (picNuA4-separated) mutant is more phenotypically similar to *Atepl1-2* than to *Ateaf1-1*.

**Figure 2.**
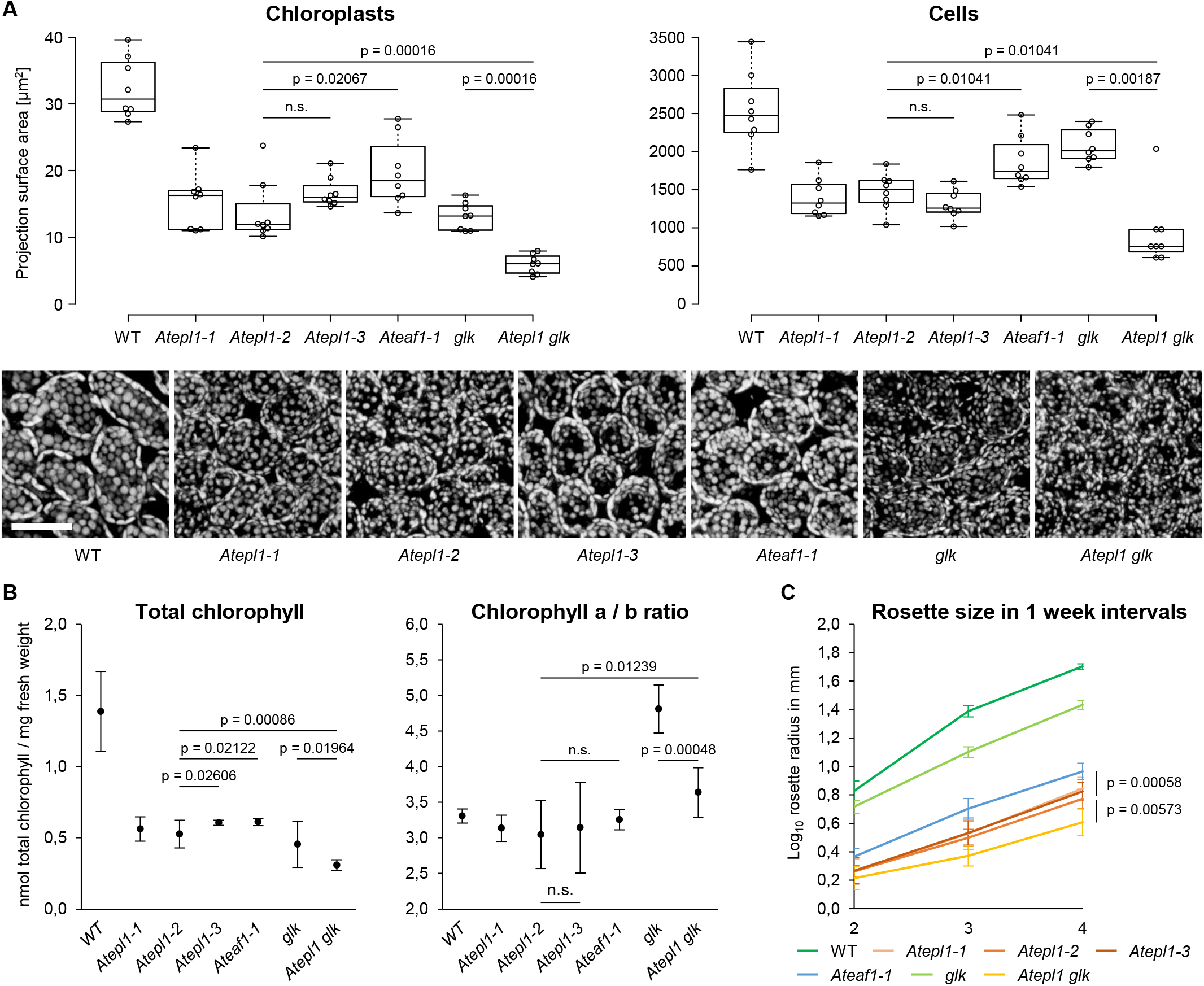
Phenotypic analysis of the NuA4 and GLK mutant lines. A. Quantification of the chloroplast- and cell-size data acquired by microscopic imaging of the palisade mesophyll cell layer. Each circle represents a median surface area of 20 chloroplasts or 9 cells measured in a single biological replicate (a single confocal stack from a separate plant). Statistical significance was determined by two-tailed Mann-Whitney U test. The chlorophyll autofluorescence images below the plots are projections of example stacks used for the measurements. All images are in the same scale. The length of the scale bar is 50 μm. B. Determination of total chlorophyll content and chlorophyll a / b ratio by spectroscopic analysis of extracted chlorophyll. Dots represent means of 3 biological replicates. The whiskers delimit 95% confidence intervals. Statistical significance was determined by two-tailed t test. C. Quantification of the growth phenotype of the mutants. The nodes represent the mean rosette radius of 10 or 20 (*Atepl1-2*) plants grown in parallel. The whiskers delimit 95% confidence intervals. P values are given for *Atea1-1* vs *Atepl1-2* (upper) and *Atepl1-2* vs *Atepl1 glk* (lower) comparisons. Statistical significance was determined by two-tailed t test.

Pale green coloration of the NuA4 and *glk* mutants indicates loss of chlorophyll. Indeed, spectrophotometric analysis of chlorophyll content in 10-leaf rosettes showed significant reduction of total chlorophyll in all mutants with the strongest effect in *glk*, followed by *Atepl1-2*. The *Ateaf1-1* mutant again displayed the weakest effect (Fig 2B, Supplementary Table 1B). Interestingly, the significant increase in the chlorophyll a / b ratio reported previously in *glk*(32) does not occur in the NuA4 mutants (Fig. 2B, Supplementary Table 1B).

In order to characterize the vegetative growth phenotype of the mutants, we measured rosette radius of soil-grown plants after 2, 3 and 4 weeks of growth in long day conditions. In this assay, the *glk* mutant performed much better than the NuA4 mutants, suggesting that the dwarfism of *Atepl1* and *Ateaf1* cannot be fully explained by chloroplast disfunction (Fig. 2C, Supplementary Table 1C). All lines which contained mutations in NuA4 subunits showed severely reduced size relative to WT plants in all time points (Fig. 2C, Supplementary Table 1C). Consistent with its relatively weak effect on cell size, *Ateaf1-1* mutation had significantly smaller effect on growth than the *Atepl1-2* mutations (Fig. 2C, Supplementary Tables 1C). Altogether, these results support a model in which AtEPL1 cooperates with AtEAF1 for most, but not all, of its functionality as a picNuA4 subunit.

### NuA4 and GLKs have parallel roles in transcription

The yeast NuA4 requires transcription factors for improved targeting to specific genomic loci(13, 33). We hypothesized that the phenotypic similarity between the mutants of NuA4 subunits and GLK transcription factors was due to targeting of NuA4 by GLKs. To verify our hypothesis, we generated a quadruple mutant *Atepl1a-2 Atepl1b-2 glk1-1 glk2-1* (later *Atepl1 glk*) by crossing *Atepl1-2* sesquimutant with the previously characterized *glk1-1 glk2-1* homozygote(27). The quadruple homozygous mutant was viable but displayed significantly aggravated phenotypic effects in all phenotypic assays (Fig 1, 2, Supplementary Table 1). Notably, the increase in chlorophyll a / b ratio observed in the *glk* but not in the NuA4 mutants was significantly dampened in the *Atepl1 glk* mutant (Fig. 2B, Supplementary Table 1B). These results suggest that NuA4 and GLK have mostly independent functions in transcription of genes related to growth and photosynthesis and may even have opposite effects on some pathways.

Seeking explanation for the mutant phenotypes and genetic interactions described above, we turned to transcriptome analysis. We analyzed RNA extracted from 10-true-leaf rosettes of *Atepl1-1* and WT plants. Overall 5319 genes showed at least two-fold decrease (hypoactive genes) or increase (hyperactive genes) of the steady-state transcript level in the mutant (Supplementary Data 1). As expected, Gene Ontology analysis of the hypoactive group (2613 uniquely mapped IDs) revealed significant enrichment in genes associated with chloroplasts, photosynthesis and growth (Fig. 3A, Supplementary Data 1). Hyperactive group (2654 uniquely mapped IDs) was significantly enriched in genes functionally related to plasma membrane as well as responses to biotic and abiotic stimuli (Fig. 3A, 3B, Supplementary Data 1). Biotic stress marker genes were also induced in *Atepl1-1* seedlings, which practically rules out a possibility of their activation by deteriorating external conditions over extended growth periods (Supplementary Fig. 6). Subsequently, we compared our results with the publicly available differential expression data for the 2-week-old *glk* mutant seedlings(34) accessed through Gene Expression Omnibus (NCBI). In contrast to the *Atepl1-1* mutant, genes differentially expressed in *glk* relative to WT control were predominantly hypoactive (866 vs 30 IDs, Fig. 3B). The small group of *glk*-hyperactive genes (30 IDs) showed negligible overlap with the *Atepl1*-hyperactive gene group (Fig. 3B). The overlap between genes hypoactive in *Atepl1-1* and *glk* was also relatively small (81 IDs). Surprisingly, genes hypoactive in *Atepl1-1* showed much higher association with chloroplasts and photosynthesis than genes hypoactive in *glk* (Fig. 3B). Finally, we used previously published transcriptomic data for inducible GLK-overexpressor lines(32) to compare *Atepl1*-hypoactive genes with *glk*-hypoactive and GLK-dependent genes (Fig. 3C). Upon closer examination of four manually curated groups of functionally related genes, we found that both NuA4 and GLKs contribute to expression of nuclear genes encoding proteins involved in chlorophyll biosynthesis and light harvesting, whereas expression of nuclear plastid ribosomal protein genes (npRPGs) depends only on NuA4 (Fig. 3C). This comparison shows that NuA4 has a much broader impact on expression of photosynthesis-related genes than GLKs (Fig. 3C). Overall, these observations are in agreement with our genetic analyses suggesting at least partially independent roles for NuA4 and GLKs in chloroplast development and maintenance.

**Figure 3.**
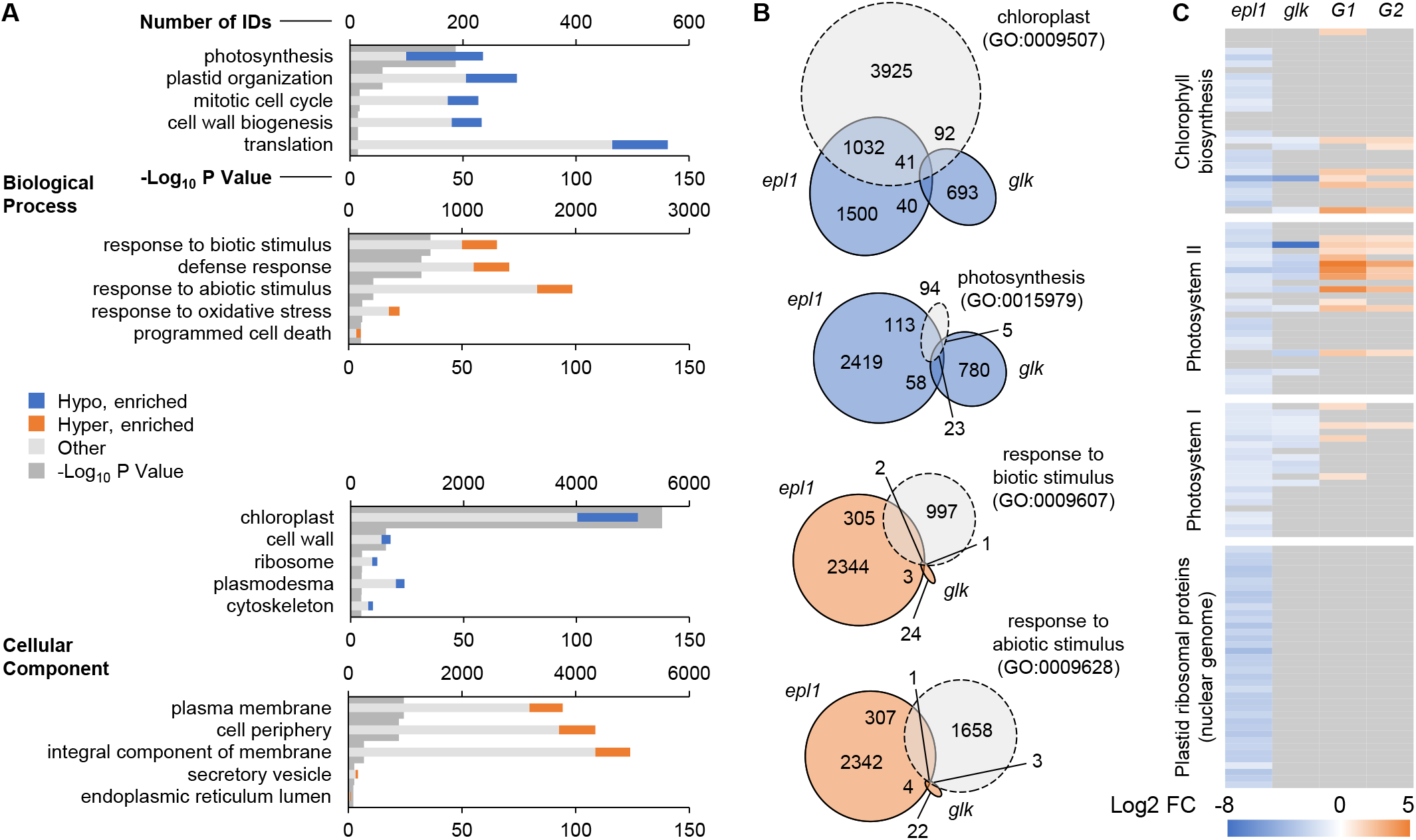
Transcriptomic analysis of the NuA4 and GLK mutants. A. Gene ontology enrichment analysis of the *Atepl1-1* hypo- and hyperactive genes. Five representative GO categories displayed in each graph were selected manually from the lists presented in Supplementary Data 1. Gene enrichment analyses were performed using PANTHER Overrepresentation Test (Supplementary Data 1). B. Area proportional Venn diagrams illustrating overlaps between genes exhibiting the same direction of expression change in *Atepl1-1* and *glk1* mutants as well as the gene sets representing particular GO categories. All numerical values indicate numbers of genes present in a single sector of the diagram rather than total values for sets. C. Relative transcript levels of genes encoding four selected groups of chloroplast proteins in *Atepl1-1*, *glk*, *GLK1ox* (*G1*) and *GLK2ox* (*G2*) lines. Expression data for the *glk* mutant and *GLKox* lines come from published works. Grey fields indicate no significant change.

### AtEPL1 is a subunit of the plant NuA4’s catalytic module

In order to further explore the molecular basis of the observed phenotypes of plant NuA4 mutants, we first had to test whether the physical interactions of AtEPL1 and AtEAF1 follow predictions based on AA sequence homology to yeast proteins. In yeast, EPL1 interacts directly with ESA1, the catalytic subunit of the complex. Arabidopsis MYST-family HATs, HAM1 and HAM2, are widely accepted as ESA1 orthologs. We produced recombinant AtEPL1A, AtEPL1B, HAM1 and HAM2 in bacteria and used two different *in vitro* methods to test direct interactions between them. Firstly, we performed a pull-down experiment in which HAM1 or HAM2 fused with His-tag were immobilized on a nickel resin and incubated with AtEPL1A or AtEPL1B fused with GST-tag. Both EPL1 homologues were specifically pulled down by both ESA1 homologues as no GST signal was detected in the eluate after His-GFP was used as a bait or when no bait was used (Fig. 4A). As a second method, we employed Far Western Blot (FWB) in which one of the recombinant proteins undergoes SDS-PAGE and Western Blot. After renaturation, the bait protein attached to the PVDF membrane is incubated with the potential interactor. Overlap between the bait and prey signal on the membrane after immunodetection of the prey indicates an interaction. Through this approach we confirmed all interactions detected by pull-down, which indicates that AtEPL1A and AtEPL1B interact directly and interchangeably with HAM1 and HAM2 (Fig. 4B).

**Figure 4.**
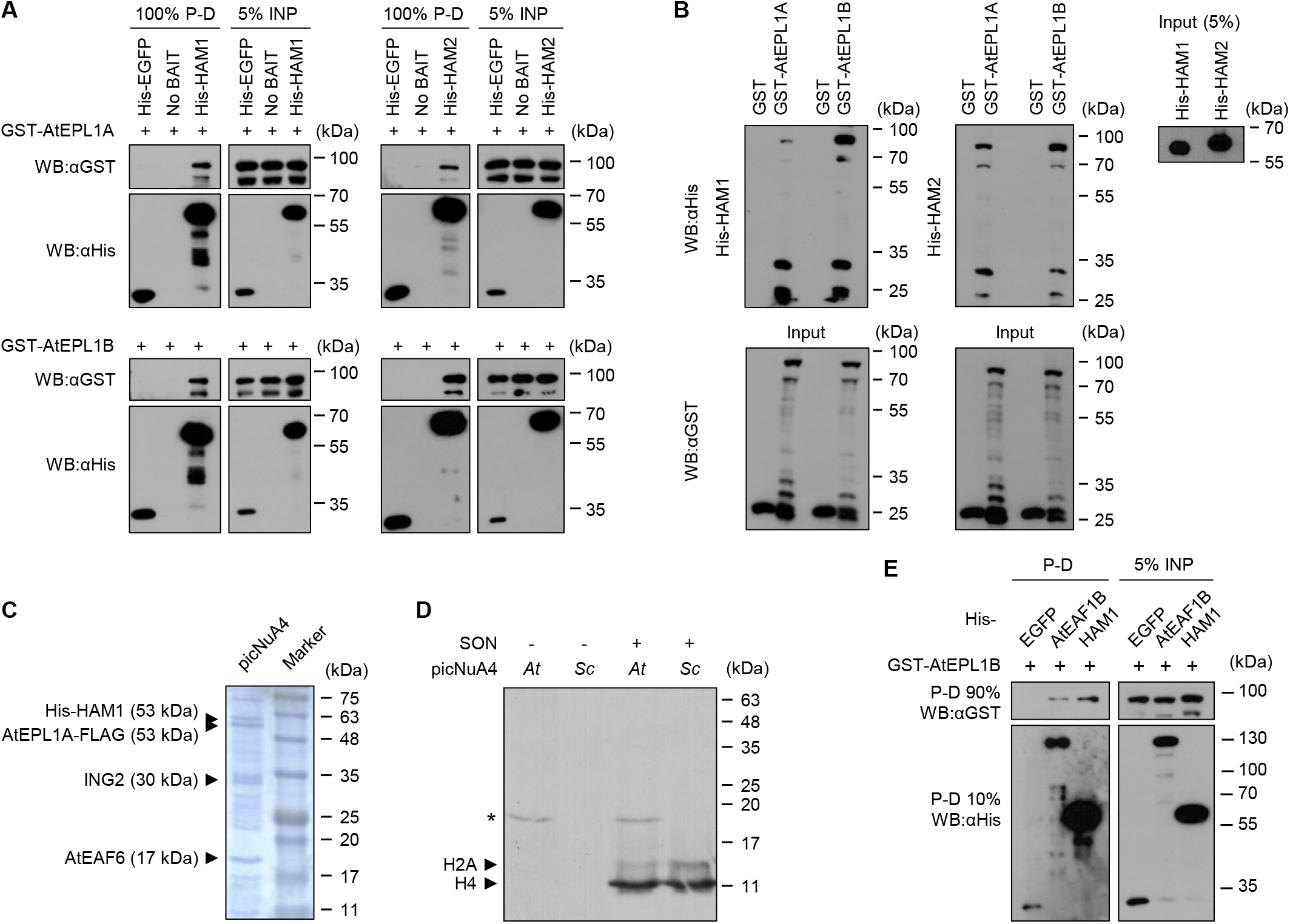
Biochemical characterization of the plant picNuA4. A. Pull-down assay showing interaction between recombinant AtEPL1 and HAM paralogue pairs expressed in bacteria. Pull-down and Input are abbreviated as P-D and INP, respectively. B. Interactions between AtEPL1s and HAMs tested by Far Western Blot approach. After SDS-PAGE and Western Blot of free GST and GST-AtEPL1A/B, the upper membranes were sequentially incubated with HisHAM1/2 and anti-His antibody whereas the lower membranes were incubated with anti-GST antibody. The lower bands in the upper panels most probably indicate HAM1/2 interaction with partially degraded AtEPL1B. C. Purification of the recombinant *A. thaliana* picNuA4 from bacteria (left panel, Coomassie staining). 6xHis-tag fused to HAM1 was used for purification on cobalt affinity resin. Numbers in brackets indicate expected molecular weights of the recombinant picNuA4 subunits. D. In vitro test of HAT activity of the recombinant *A. thaliana* (*At*) picNuA4 towards short oligonucleosomes (SON). Recombinant *S. cerevisiae* (*Sc*) picNuA4 was used as a positive control. Autoacetylation on AtEAF6 is highlighted by an asterisk. E. Pull-down assay showing the interaction between AtEPL1B and a fragment of AtEAF1B corresponding to AA 430-1322. HAM1 is used for comparison. All proteins were expressed in bacteria as either GST- or 6xHis-fusions.

In yeast, the functional significance of the physical interaction between EPL1 and ESA1 lies in the ability of picNuA4 (formed through binding of ESA1, YNG2 and EAF6 to EPL1) to acetylate histones in the context of a nucleosome. In order to test if *A. thaliana* picNuA4 is capable of nucleosomal histone acetylation, we co-expressed HAM1, AtEPL1A, ING2 and AtEAF6 in bacteria and used the recombinant complex for in vitro acetylation assay (Fig. 4C, 4D). Notably, using HAM1 as a bait for complex purification resulted in recovery of all recombinant picNuA4 subunits which confirms that these four plant proteins interact physically to form a complex homologous to yeast picNuA4 (Fig. 4C). The *in vitro* HAT assay confirmed the activity of the Arabidopsis picNuA4 on native oligonucleosomes. As expected, tritium-labeled acetyl groups were efficiently transferred to histones H2A and H4, with some preference for the latter, when compared to the yeast complex (Fig. 4D). These results provide strong evidence for evolutionary conservation of the subunit composition and the activity of the picNuA4 complex between budding yeast and plants.

### AtEPL1 interacts with the subunits of the NuA4’s regulatory module

PicNuA4 interacts with the targeting module of the yeast NuA4 through the contact made by the EPcB domain of EPL1 with EAF1. Our phenotypic analyses of the *Atepl1-3* and *Ateaf1-1* mutants suggest that AtEAF1 and the EPcB domain of AtEPL1 are crucial elements of the plant NuA4 complex. In order to find biochemical confirmation of this observation we performed tandem affinity purification followed by mass spectrometry in which we used the *Atepl1-1* mutant complemented with the *pAtEPL1B∷AtEPL1B-TAP* construct. The experiment was performed twice on hydroponically grown seedling. Notably, out of the 15 proteins specifically copurified with AtEPL1B in both experiments, 12 were homologues of yeast NuA4 subunits, including EAF1 and ESA1 (Table 1, Supplementary Data 2). This result confirms and improves on previously published findings regarding potential protein interactors of AtEPL1B(30) and other putative subunits of *A. thaliana* NuA4(22, 24, 30, 35) (summarized in Table 1). In particular, the use of tandem affinity purification resulted in low total number of interactors but high number of NuA4 subunit-homologues, indicating high specificity (Table 1, Supplementary Data 2). In fact, our analysis identified only 5 proteins which show no homology to yeast NuA4 subunits as potential AtEPL1B interactors (Table 1, Supplementary Data 2). Two of these proteins, AOG2 and BRD6, were copurified with *A. thaliana* NuA4 subunits previously, albeit in much larger pools of additional interactors (Table 1)(24, 30, 36). Interestingly, the uncharacterized bromodomain protein BRD6(37), previously copurified with SWC4(24) and HAM1(30) (Table 1), is orthologous to BRD8, a subunit of the human equivalent of NuA4, the TIP60/p400 complex, according to PANTHER Classification System(38, 39), indicating a possible role as an accessory subunit of the plant NuA4. AOG2, on the other hand, may be of interest in the context of the role of NuA4 in gametophyte development(36).

**Table 1.**
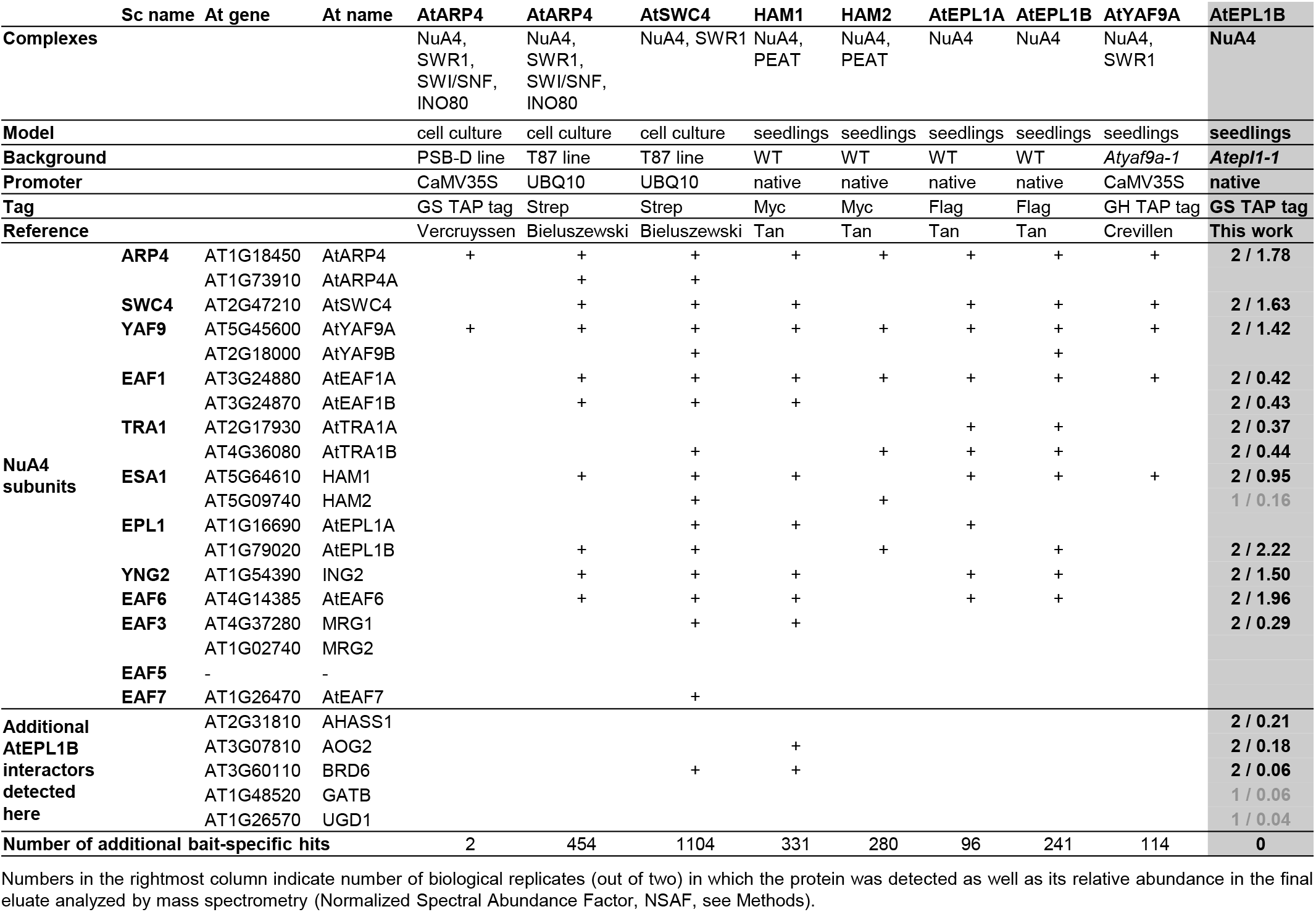
TAP-MS data for AtEPL1B combined with metaanalysis of the previously published AP-MS experiments using NuA4 subunits as baits.

In order to test whether AtEPL1 interacts with the targeting part of NuA4 through direct binding to AtEAF1, we performed pull-down assay in which a His-tagged fragment of AtEAF1B corresponding to AAs 430-1322, His-HAM1 (positive control) or His-EGFP (negative control) were used as baits and GST-AtEPL1B was used as prey. Only His-AtEAF1B and His-HAM1 were able to pull-down GST-EPL1B which confirmed a direct physical interaction between AtEPL1B and AtEAF1B (Fig. 4E). These results support our homology-based model in which AtEPL1 and AtEAF1 interact directly to provide a scaffold for the assembly of the plant NuA4 complex.

### Loss of AtEPL1 triggers genome-wide depletion of H4K5ac but not H3K9ac

Despite global role of NuA4 in maintaining H4 acetylation, studies in yeast suggest stronger NuA4-dependence in certain groups of genes such as genes related to ribosome biogenesis(18, 40). We hypothesized that expression of some nuclear genome-encoded chloroplast- and photosynthesis-related genes may depend on acetylation of H4K5 by plant NuA4. Previous studies showed that expression of light-regulated genes in plants correlates with and depends on local histone acetylation status, especially that of H3K9(8, 9). In order to test our hypothesis, we profiled the NuA4-dependent H4K5ac and the GCN5/SAGA-dependent H3K9ac genome-wide in the *Atepl1-2* mutant and WT plants. We expected some genes hypoactive in the mutant to show reduced H3K9ac due to indirect effects such as retrograde signaling. Conversely, preservation of H3K9ac coinciding with highly reduced H4K5ac and transcript levels would point to direct NuA4-dependence.

We performed Chromatin Immunoprecipitation followed by DNA-sequencing (ChIP-seq) on 10 true-leaf rosettes of *Atepl1-2* and WT grown in the same conditions as plants used for RNA-seq. In addition to H4K5ac and H3K9ac, we used antibodies specific to unmodified H3 for normalization of the acetyl-mark signal to nucleosome occupancy. Due to expected global changes in epitope abundance, we included a predetermined proportion of mouse chromatin in our ChIPs as a reference (“spike-in”).

As expected, NuA4 inactivation in the *Atepl1-2* mutant resulted in globally reduced levels of H4K5ac mark. Its occupancy profiles, averaged across all expressed genes, show strong reduction in the mutant throughout the 3000 bp region centered on TSS (Fig. 5A, Supplementary Fig. 8). In contrast to H4K5ac, the profile of H3K9ac in *Atepl1-2* overlaps well with WT, with only a slight increase within the peak downstream of TSS. Interestingly, the averaged H3 occupancy in *Atepl1-2* showed a marked increase on both sides of TSS (Fig. 5A, Supplementary Fig. 8).

**Figure 5.**
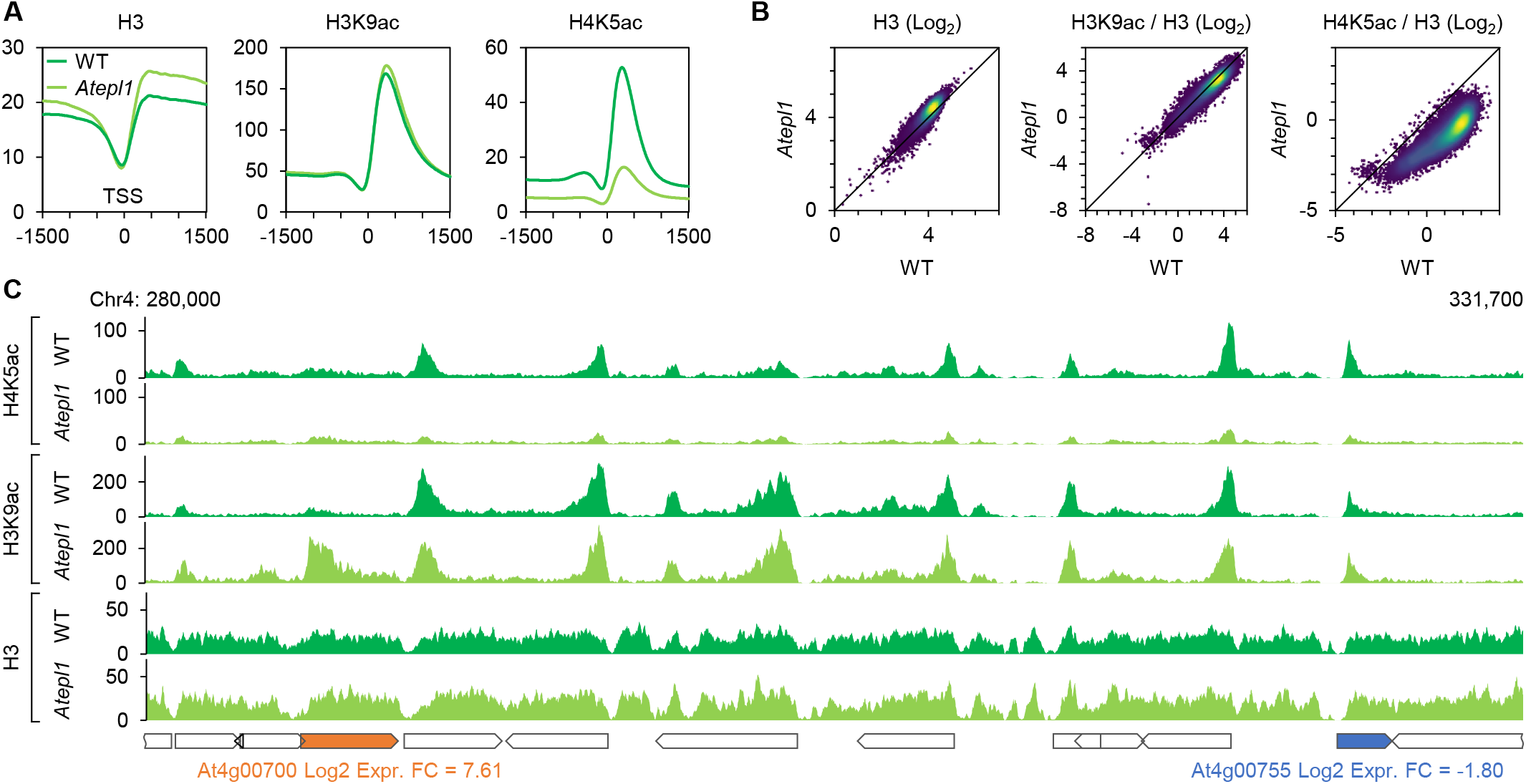
Chromatin landscape of the *Atepl1-2* mutant. A. TSS-centered profiles of spike-in-normalized occupancy of H3, H3K9ac or H4K5ac averaged over all genes expressed in WT Col-0. Vertical axes values are relative. Horizontal axes values represent chromosomal position relative to TSS (i.e. negative values represent promoter region whereas positive values represent gene body). B. Scatter plots showing differences in H3, H3K9ac or H4K5ac occupancy between *Atepl1-2* and WT. Each dot represents Log_2_-transformed ChIP-seq signal averaged over the first 500 bp downstream from TSS of a single gene in WT and in *Atepl1-2*. The number of dots per pixel is color-coded. C. A fragment of chromosome 4 showing spike-in-normalized H3, H3K9ac or H4K5ac occupancy across several loci. The particular chromosomal location was chosen to illustrate chromatin environment changes coinciding with increased (orange gene) or decreased (blue) transcript levels versus no significant change (white). Vertical axes values are relative.

As both acetyl marks are highly enriched downstream of TSS in the WT (Fig. 5A), we averaged the signal over the first 500 bp of each transcribed region and used these values to make comparisons between individual genes. While H3-normalized H4K5ac levels in *Atepl1-2* showed strong and almost uniform reduction in this window, H3K9ac levels were preserved in most genes, with few genes showing strong changes (Fig. 5B). In agreement with the role of H3K9ac in transcriptional activation, we observed increased H3K9ac levels over strongly *Atepl1*-hyperactive genes (Fig. 5C). On the other hand, despite strong H4K5ac reduction over most genes, only some from them showed reduced transcript levels (Fig. 5C) suggesting that NuA4-dependency is determined by additional locus-specific factors.

### Expression of npRPGs specifically depends on H4 acetylation by NuA4

Due to synergistic action of NuA4 and SAGA over most genes in yeast(18), one plausible explanation of decreased expression of some genes is a concomitant loss of both acetylation marks in *Atepl1* mutants. In such cases, there is no basis for a direct link between their expression and NuA4 activity. On the other hand, genes hypoactive in *Atepl1-2* showing no significant reduction in H3K9ac but strong reduction in H4K5ac are potentially directly affected by loss of H4K5ac. In order to strictly define this group of genes we set the following arbitrary criteria: (i) Log_2_ of transcript level fold change should be lower than −1.5, (ii) Log_2_ of H4K5ac fold change should be lower than −2, (iii) Log_2_ of H3K9ac fold change should be higher than −0.5 (Fig. 6A, Supplementary Data 3). We visualized distribution of all genes hypo- or hyperactive in *Atepl1-1* across these three dimensions by plotting log_2_-transformed fold change values of the two acetyl-marks against each other, separately for hypo- and hyperactive genes. (Fig. 6B). Notably, *Atepl1*-hyperactive genes are shifted towards the upper right corner of the plot (increased H3K9ac, no reduction in H4K5ac), indicating a role for H3K9ac in their increased activity in the mutant as well as some level of dependency of H4K5ac on H3K9ac or transcription in *Atepl1* background. Distribution of the hypoactive genes is less skewed along the Y-axis (i.e. roughly equal numbers of hypoactive genes gain and lose H3K9ac) which suggests that their behavior is driven mostly by loss of H4K5ac. Application of the acetylation fold change criteria (ii) and (iii) to *Atepl1*-hypoactive genes leaves 350 potentially NuA4-dependent genes (Supplementary Data 3). Intriguingly, when tested against all *Atepl1*-hypoactive genes, this set is significantly enriched in genes associated with growth and photosynthesis (Fig. 6C, Supplementary Data 3). Nuclear genes encoding plastid ribosomal proteins(28) deserve particular attention as nearly all of them cluster within or near the NuA4-dependent sector (Fig. 6B, Supplementary Data 3). Furthermore, the median log_2_ expression FC of translation- and ribosome-related NuA4-dependent genes is significantly lower than that of all hypoactive genes as could be expected from genes which directly depend on NuA4 for efficient transcription (Fig. 6D). This suggests that NuA4 complex has been adopted to specifically stimulate expression of growth and photosynthesis-related genes in plants.

**Figure 6.**
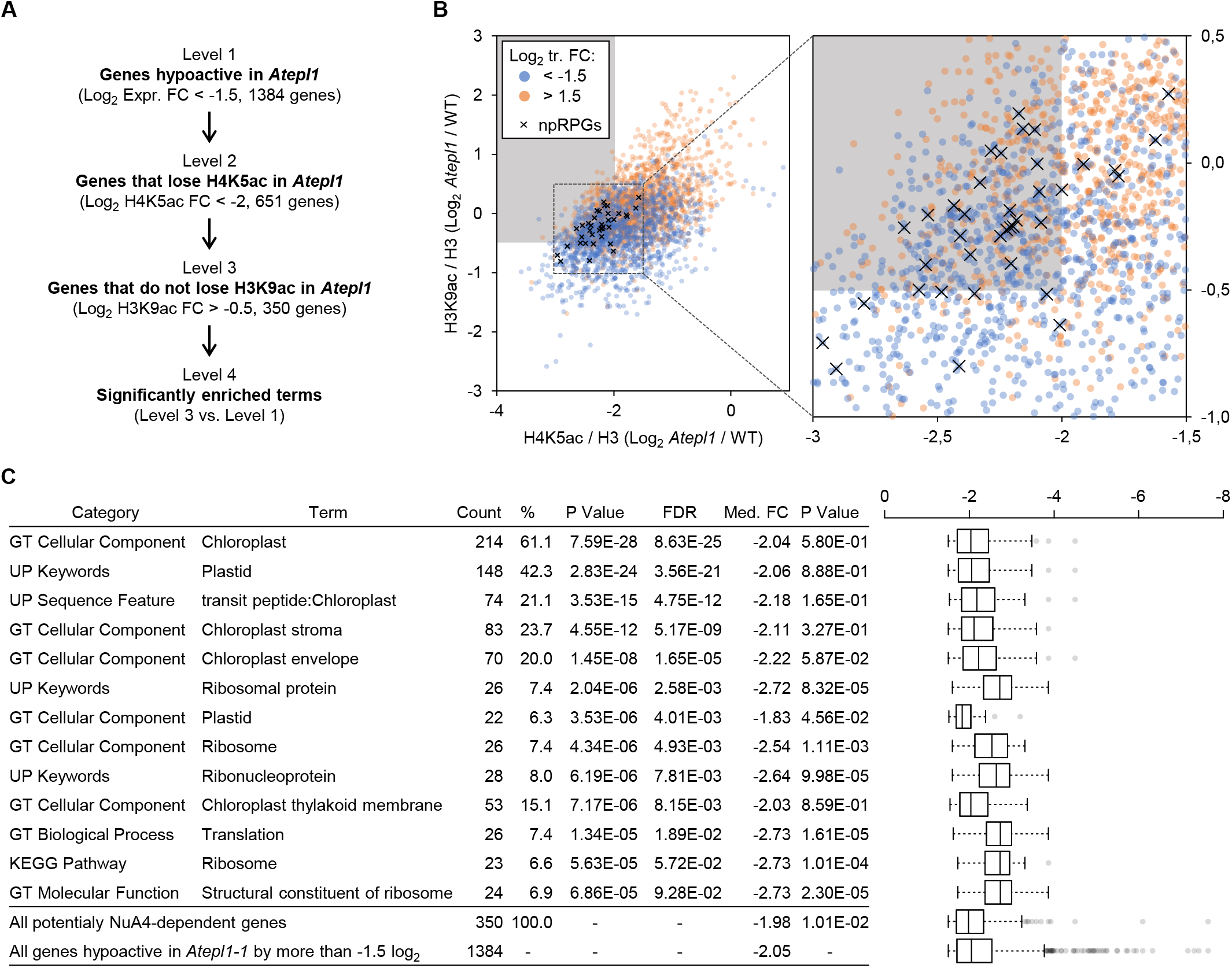
Identification of the potentially NuA4-dependent genes. A. Ideogram of the NuA4-dependency determination process. B. Scatter plot showing relationships between Log_2_-transformed fold-change of the spike-in- and H3-normalized H3K9ac and H4K5ac occupancy and the directionality of the transcript level change. The dashed line delimits the “NuA4-dependent” area of the plot as described in (A). C. Functional enrichment and relative expression of NuA4-dependent genes. P value for functional enrichment (column 5) is equivalent to EASE score (modified Fisher Exact test, see Methods). P values for the differences between median fold change against all genes hypoactive in *Atepl1-1* (column 8) are results of Mann Whitney U test. GT stands for GO Term.

## Discussion

Our previous attempts at methodical genetic analysis of the plant NuA4 complex were hindered by the tandem duplication of the gene encoding *A. thaliana* homologue of budding yeast EAF1(24). The Cas9-induced loss of function mutant presented here brings new evidence supporting a central role of AtEAF1 in the plant NuA4 (Fig. 1, 2). First of all, the phenotype of the *Ateaf1-1* mutant is significantly more severe than the phenotypes of the *mrg*, *eaf7* and *yaf9* mutants described previously(20, 22, 23). We note that lethal effects have been reported for the *Atswc4* mutant as well as the double *ham* mutant(19, 41). These are probably not NuA4-specific, however, due to AtSWC4’s role in the SWR1 complex and potential NuA4-independent roles of HAM in acetylation of non-histone substrates(41, 42). Furthermore, the *Ateaf1-1* mutant displays an extreme enhancement of the photosynthetic and reproductive phenotypes previously reported in the double mutant of AtYAF9, a confirmed interactor of AtEAF1(22, 24).

In the absence of EAF1, picNuA4 remains capable of non-targeted nucleosomal histone acetylation, enabling survival(14, 43). Deletion of the picNuA4-specific subunit EPL1/EPC1 is lethal in yeast and in animals(44, 45). Our biochemical analyses of the two previously identified *A. thaliana* homologues of EPL1(24, 46) link it unequivocally to the plant NuA4 complex and acetylation of nucleosomal substrates (Fig. 4). Remarkably, we show that plants tolerate loss of EPL1 despite dramatic genome-wide reduction of H4K5ac and severe disorganization of the transcriptome (Fig. 1, 2, 3, 5). This unexpected result should be confronted with a recent study showing that, in addition to the pair of EPL1 orthologues described herein, *A. thaliana* contains two EPL1-related proteins (EPCR)(30). Despite marginal sequence similarity to EPL1, EPCRs interact physically with HAM1 and HAM2(30), possibly explaining why essential functions of HAMs and picNuA4 may be completed without AtEPL1. On the other hand, EPCRs do not copurify with any other subunit of the plant NuA4(30) and the HAT activity of the EPCR-HAM complex has not been proven yet, therefore, this explanation is only hypothetical.

Severe phenotype of *Ateaf1-1*, most probably caused by dissolution of the regulatory module of NuA4, suggests that proper targeting of the picNuA4 activity is outstandingly important in plants. To our knowledge, the phenotype of *Ateaf1-1* is not copied by any mutation disrupting a nuclear transcription factor described to date in *A. thaliana*. Nevertheless, we found the cell-autonomous photosynthetic phenotype shared by *Ateaf1* and *Atepl1* mutants to be reminiscent of the well characterized double *glk* mutant (Fig. 2)(27, 31). We hypothesized that GLKs guide NuA4 to genes related to chloroplast development and photosynthesis. Our genetic experiments and transcriptomic analyses provide convincing evidence against this hypothesis. Although *Ateaf1*, *Atepl1* and *glk* mutants display comparable levels of reduction in chloroplast size and chlorophyll level, these phenotypes are enhanced in the quadruple *Atepl1 glk* mutant, suggesting at least partial functional independence (Fig. 1, 2). While GLKs directly control transcript levels of a relatively small set of genes, NuA4 influences a significant proportion of all genes related to chloroplasts and photosynthesis with a particularly strong effect on GLK-independent npRPGs (Fig. 3). Finally, GLKs were not copurified here with EPL1B nor with any other NuA4 subunit used as a bait in the AP-MS experiments reported to date (Table 1). All this implies that NuA4 and GLK operate mostly independently although a limited cooperation in some loci cannot be excluded.

Since phenotypes of *Ateaf1* and *Atepl1* mutants differed quantitatively rather than qualitatively (Fig. 1, 2), we assumed similarly widespread changes in H4K5ac occupancy in both cases. Consequently, we focused on the *Atepl1-2* mutant which gave us a unique opportunity to investigate the impact of complete NuA4 inactivation while at the same time providing hints to explain the common phenotype. Interestingly, despite roughly equal numbers of hypo- and hyper-active genes in *Atepl1*, the H4K5ac levels were globally decreased (Fig. 3, 5). This suggests either synergistic control by various acetyl marks or no influence of H4K5ac on transcription in most loci. Another important observation is the lack of a global negative influence of H4K5ac reduction on H3K9ac levels. Such influence has been recently observed in ESA1-depleted budding yeast(18). Furthermore, many of the *Atepl1*-hyperactive genes became hyperacetylated on H3K9 in the mutant despite showing low acetylation levels in the WT (Fig. 5, 6), suggesting that H3K9ac may effectively build up and stimulate transcription in the absence of nucleosomal H4K5 acetylation in plants.

In yeast, certain groups of genes show high NuA4-dependence. These are mostly genes involved in translation and other anabolic processes(18, 33, 40, 47, 48). Consistently, NuA4-defficient yeast strains are hypersensitive to rapamycin which inhibits growth-promoting TOR signaling(14). Our results suggest that in plants NuA4-dependence is shifted from cytoplasmic to plastidic translation machinery. To the best of our knowledge none of chromatin factors involved in transcription control characterized to date show similar photosynthetic-related phenotypes while inactivated in plants. The nucleus-encoded plastid ribosomal protein genes showed little loss of H3K9ac but strong and uniform reduction in H4K5ac and transcript levels, indicating potential direct NuA4-dependence (Fig. 7). Since these and other nuclear genes encoding chloroplastic proteins originated from a cyanobacterial endosymbiont, there is an ongoing search for a mechanism which enabled their effective transfer to the nucleus. It has been proposed that open chromatin configuration facilitates both acquisition and transcriptional activity of new mitochondrial and chloroplastic insertions in the nuclear genome(49, 50). It is not unlikely that further investigation of the mechanisms through which NuA4 affects chromatin structure and transcription in plants will improve our understanding of the transition between the endosymbiont and the chloroplast at the origin of the green lineage.

## Materials and methods

### Plant material

All *A. thaliana* mutants used in this work are in Col-0 background. T-DNA insertion mutants for AtEPL1A (At1g16690, SAIL_239_D12(51), *Atepl1a-1*) and AtEPL1B (At1g79020, SALK_094941(52), *Atepl1b-1*) were obtained from NASC and crossed to produce the double mutant *Atepl1-1*. The previously characterized double homozygous transposon-insertion mutant *glk1-1 glk2-1*(27) also comes from the NASC collection (NASC ID: N9807). All other mutants were generated for this work through Cas9-gRNA mediated genome editing of WT Col-0 plants and are described in detail in Supplementary Figures 1-3.

### Mutant generation through Cas9-gRNA

#### Preparation of genetic constructs

The Cas9 expression cassette was created through fusion of the *ICU2* promoter sequence cloned from genomic DNA following published instructions(53) with the 3xFlag-NLS-hSpCas9-NLS∷NosT cassette which was a gift from Dr Jian-Kang Zhu(54). Following insertion of the Cas9 expression cassette into the pFGC vector (a gift of Dr Jian-Feng Li(55)), the *Bam*HI site was moved from the 5’- to the 3’-end of the Nos terminator. The *Bam*HI site was subsequently used for linearization of the plasmid and insertion of gRNA cassettes by Gibson assembly-based cloning. The gRNA expression cassette containing U6 promoter was a gift from Dr Jian-Kang Zhu(54). The U3 expression cassette was created by replacing the U6 promoter with the *AtU3C* promoter cloned from genomic DNA isolated from *A. thaliana* Ler ecotype following published sequence data(56). The target sequences were found with CRISPR-P(57) (*Ateaf1-1*, *Atepl1-2*) or CRISPOR(58) (*Atepl1-3*). Targeted mutagenesis of the gRNA expression cassettes was done by PCR-amplification and recircularization of the pJET1.2-based gRNA vectors. The sequences of the primers used for mutagenesis can be found in the Supplementary Table 3. **Editing strategy and isolation of the mutants.** In order to generate genomic deletions within coding regions of the target genes, double-gRNA-Cas9 constructs were integrated with the genome through Agrobacterium mediated transformation(59). Transformed plants were selected on soil by spraying with Basta (Bayer). Two primers flanking the expected deletion were used to identify plants which accumulated somatic mutations apparent from the presence of short PCR products corresponding to deletions between two neighboring cut sites. T2 plants were screened for the absence of Cas9-gRNA T-DNA by PCR and genotyped again for the presence of desired genomic deletions. Any mutations detected in this way were assumed heritable. Subsequently PCR amplification products were cloned into pJET1.2 vector and plasmids isolated from multiple single colonies were sequenced to determine the sequences of both homologous loci. If plants were confirmed as frame-shift- (*Ateaf1-1*, *Atepl1-2*) or in-frame-deletion mutants (*Atepl1-3*), they were backcrossed to Col-0 and propagated. Sequences of all primers used for genotyping can be found in Supplementary Table 3.

### Phenotypic analyses

For all phenotypic analyses plants were grown in peat pellets (Jiffy) in controlled conditions: 22°C, 80-90 μmol/(m^2^s) PAR, 60% RH. In course of the growth rate analyses, plants were imaged in one-week intervals with a stereomicroscope or a camera, depending on the size. ImageJ software was used for image analyses to extract the distance between the tip of the longest rosette leaf and rosette center(60).

For plastid- and cell-size measurements, leaves #4 were cut from 10-true-leaf rosettes, vacuum-infiltrated with perfluoroperhydrophenanthrene(61), mounted on a slide and imaged with the A1Rsi confocal microscope (Nikon). Z-stacks spanning the palisade mesophyll cell-layer, acquired in transmitted-light and chlorophyll autofluorescence channels, were used for analysis in ImageJ. Contours of cells and plastids were tracked manually and the surface area of the resulting regions of interest (ROI) was treated as the surface projection area of the measured structures. All calculations and statistical analysis was done using Microsoft Excel.

Chlorophyll was extracted from a single plant for each biological replicate. A 10-leaf rosette was separated from the root, weighed and grinded in liquid nitrogen. After obtaining a fine powder, 3 or 5 ml (depending on rosette size) of Tris-HCl-buffered 80% acetone was added and grinding was continued until homogeneity. The thawed mixture was then transferred to a 10 ml glass tube wrapped in aluminum foil to prevent access of light, sealed and kept in 4°C overnight. Directly before spectroscopic measurements 1.5 ml of the mixture was transferred to 1.5 ml microtubes (Axygen) and centrifuged at 10,000 G for 5 min to pellet any debris. After centrifugation the extract was transferred to a spectrophotometer cuvette and absorbance was measured at 647 nm and 664 nm using BioSpectrometer kinetic (Eppendorf) spectrophotometer against buffered acetone as blank. The number of samples processed in parallel was limited to 4 in order to reduce the influence of the processing time on the measurements. Total chlorophyll content as well as chlorophyll a / b ratio were calculated according to published formulas(62) without correction for absorbance at 750 nm.

### Pull-down

Recombinant proteins were expressed in *E. coli* strains BL21-CodonPlus-RP or Lemo21(DE3, New England Biolabs). Bacterial lysates were prepared as previously described(48) with the following modifications. GST-tag and His-tag fusion proteins were resuspended in PBS or equilibration and wash buffer (50 mM sodium phosphate, pH 8.0, with 0.3 M sodium chloride, 10 mM imidazole) respectively, with addition of 1 mM phenylmethylsulfonyl fluoride (PMSF) and cOmplete™ Mini EDTA-free Protease Inhibitor Cocktail (Roche). His-GFP or His-fusion proteins were bound to HIS-Selec HF Nickel Affinity Gel (Sigma) for 1 hour, then washed and incubated with recombinant GST-fusion proteins for additional 1 h at 4°C. The beads were washed five times with wash buffer containing 0,5% NP40. The protein complexes were analyzed by SDS/PAGE followed by WB analysis with anti-GST (B14; #sc-138; Santa Cruz) and anti-His (#R940-25; Life Technology, Waltham, MA, USA) antibodies.

### Far Western Blotting

This experiment was performed as previously described(63) with some modification. Briefly, GST, GST-AtEPL1A and GST-AtEPL1B were purified on Glutathione Sepharose 4 Fast Flow beads (GE Healthcare) for 1 h at 4°C. The beads were washed five times with PBS containing 1% Triton X-100. The GST-fusion proteins were eluted by heating at 95°C in SDS/PAGE loading buffer, resolved by SDS/PAGE and transferred onto PVDF membrane. Proteins were renatured by incubation of the membrane in AC buffer (20 mM Tris pH 7.5, 1 mM EDTA, 0.1 M NaCl, 10% glycerol, 0.1% Tween-20, 2% non-fat milk, 1 mM DTT) containing 6 M guanidine-HCl for 30 min at RT. Subsequently, the membrane was washed with AC buffer containing 3 M guanidine-HCl for 30 min at RT. This step was followed by washing with AC buffer containing 0.1 M- and no-guanidine-HCl AC buffer at 4°C, for 30 min and overnight respectively. The membrane was blocked for 8 h at 4 °C in 5% nonfat milk in the PBST buffer and subsequently incubated with His-HAM1 or His-HAM2. Next, the membrane was washed with 5% nonfat milk in the PBST buffer followed by three washes with PBST. To detect bait proteins bound to the bait proteins (GST-fusions) on the membrane, standard immunodetection with anti-His antibody (#R940-25; Life Technology, Waltham, MA, USA) was performed.

### In vitro acetylation assay

#### Genetic construct preparation

Coding sequences of AtEPL1A, HAM1, ING2 and AtEAF6 were cloned to the pST44 polycistronic vector for protein expression in *E. coli*, according to the original pST44 documentation(64). FLAG-tag was added to the N-terminus of AtEPL1A and 6xHis-tag was added to the C-terminus of HAM1. **Protein expression and purification.** Protein expression was conducted in BL21(DE3)pLysS cells after 0.3 mM IPTG induction and the complex was purified over Talon (Clontech) cobalt affinity resin as described previously(65) via Ham1-6xHis, using 150 mM imidazole for elution. Presence of all subunits was visible on Coomassie-stained SDS-PA gel and presence of HAM1-His and FLAG-AtEPL1A was confirmed by Western blot. **HAT assays.** HAT assays were performed as described previously(66) with some changes. Briefly, in liquid HAT assays 0.5 μg of purified human core histones or native H1-depleted short oligonucleosomes were incubated together with bacterially expressed plant picNuA4 (or yeast picNuA4 as a positive control) complexes with ^3^H-labeled acetyl-CoA (0.125 μCi) in HAT buffer (50 mM Tris-HCl pH 8.0, 50 mM [KCl+NaCl], 5% glycerol, 0.1 mM EDTA, 1 mM DTT, 1 mM PMSF, 10 mM sodium butyrate) at 30°C for 30 min. Subsequently, the reactions were spotted on P81 membrane (Whatman), which were then air-dried, washed 3 times with 50 mM carbonate buffer pH 9.2, rinsed in acetone and after soaking in scintillation cocktail used to count ^3^H signal. Alternatively, reactions were resolved by SDS-PAGE. After electrophoresis gel was stained in Coomassie to visualize proteins, destained and incubated in EN3HANCE solution (Perkin Elmer). After washing and drying, gel was incubated with film.

### Tandem Affinity Purification (TAP)

The construct used for purification of proteins interacting with AtEPL1B was created by replacing the Gateway cassette in pMDC99 vector with the *AtEPL1B* expression cassette. The *AtEPL1B* cassette consisting of the genomic sequence of *AtEPL1B* (At1g79020) including 969 bp of the native upstream regulatory sequence, C-terminal GSrhino tag(67), native 3’ UTR of *AtEPL1B* and Nos terminator was assembled in the pMDC99 backbone through Gibson assembly-based cloning. Growing of seedlings and TAP purifications using 50 g of six-day old seedlings as input per experiment, were performed as described in (67). Bound proteins were digested on-bead after a final wash with 500 μL 50 mM NH_4_HCO_3_ (pH 8.0). Beads were incubated with 1 μg Trypsin/Lys-C in 50 μL 50 mM NH_4_OH and incubated at 37°C overnight in a thermomixer at 800 rpm. Next day, an extra 0.5 μg Trypsin/Lys-C was added and the digest was further incubated for 2 hours at 37°C. Finally, the digest was separated from the beads and centrifuged at 20800 rcf in an Eppendorf centrifuge for 5 min, the supernatant was transferred to a new 1.5 mL Eppendorf tube. Peptides were purified on C18 Bond Elut OMIX tips (Agilent) and purified peptides were dried in a Speedvac and stored at −20°C until MS analysis. Co-purified proteins were identified by mass spectrometry using a Q Exactive mass spectrometer (ThermoFisher Scientific) using standard procedures(68)). Proteins with at least two matched high confident peptides were retained. Background proteins were filtered out based on frequency of occurrence of the co-purified proteins in a large dataset containing 543 TAP experiments using 115 different baits(67). True interactors that might have been filtered out because of their presence in the list of non-specific proteins were retained by means of semi-quantitative analysis using the average normalized spectral abundance factors (NSAF) of the identified proteins in the EPL1B TAPs. Proteins identified with at least two peptides in at least two experiments, showing high (at least 10-fold) and significant [-log10(p-value(T-test)) ≥10] enrichment compared to calculated average NSAF values from a large dataset of TAP experiments with non-related bait proteins, were retained(69).

### RNA-seq

For RNA-seq WT and *Atepl1-1* plants were grown under LD conditions (16 h light and 8 h darkness), under 80-90 μmol/(m^2^s) of photosynthetically active radiation. Material was collected at the developmental stage of ca. 10 leaves about 5 hours after light onset.

RNA for each biological repeat was extracted from 110 mg of rosette leaves number 5-6 (from at least six plants) with TRI Reagent (Merck) and rounds of phenol-chloroform and chloroform extractions followed by isopropanol precipitation. RNA was treated with RQ1 DNase I (Promega), then extracted with phenol-chloroform and precipitated with ethanol. Pure RNA solution was sent to Macrogen Europe (South Korea), where libraries and sequencing were performed using Illumina chemistry. RNA-seq experiment was also validated by RT-qPCR (Supplementary Fig. 5).

Adapters, poor quality regions (mean Phred quality < 5) and rRNA sequences were removed from short reads using BBDuk2 program from BBTools suite (https://jgi.doe.gov/data-and-tools/bbtools). Filtered reads shorter than 50 bp were excluded from the analysis. Expression values for genes and transcripts (Araport11 annotations) were calculated using Salmon(70) with sequence-specific bias and GC content bias correction. Differential expression analysis on both gene and transcript level was performed using limma package(71) following a previously published protocol(72). False discovery rate (FDR) threshold of 0.01 and fold change threshold of Log_2_FC=1 (Fig. 3) or Log_2_FC=1.5 (Fig. 6) were used in the analyses.

### ChIP-seq

ChIP-seq analysis was carried out on WT and *Atepl1-2* plants grown in the same conditions as the plants used for RNA-seq and harvested at the same developmental stage. Chromatin was isolated as previously described(73), enriched with 2% chromatin isolated from mouse embryonic fibroblasts as spike-in control, and incubated with antibody (Abcam ab1791, Millipore 07327, Millipore 07352) overnight at 4°C. Dynabeads Protein A (Thermo Fisher Scientific) were added to the samples and incubated for another 2 hours at 4°C. The slurry was washed twice with low salt buffer (with 150 mM NaCl), twice with high salt buffer (with 500 mM NaCl) and once with LiCl wash buffer (10 mM Tris-HCl, pH 8.0, 1 mM EDTA, 0.25 M LiCl, 1% Nonidet P-40, 0.5% sodium deoxycholate, 1 mM PMSF and protease inhibitor cocktail). Chromatin was eluted through incubation in 1% SDS and 0.8 M NaHCO3 at 65°C and the cross-links were reversed by overnight incubation at 65°C. Following Proteinase K treatment and phenol/chloroform extraction, DNA was precipitated and DNA concentration was measured with Qubit dsDNA HS Kit (Thermo Fisher Scientific). 10 ng (H3) or 2 ng (other antibodies) of ChIP-ed DNA was used to prepare DNA libraries with MicroPlex Library Preparation Kit v2 (Diagenode).

Adapters and poor-quality regions (mean Phred quality < 5) were removed from short reads using BBDuk2 program from BBTools suite (https://jgi.doe.gov/data-and-tools/bbtools). Filtered reads shorter than 35 bp were excluded from the analysis. Reads were then aligned to the *A. thaliana* (TAIR10) and mouse (GRCm38) genomes using BWA-MEM aligner(74). Sambamba(75) was used to process the aligned data and remove duplicated reads. Mapping statistics, including the percentages of mapped reads and duplicates, were calculated separately for *A. thaliana* genome (ChIP-Seq data) and mouse genome (“spike-in” data) using BamTools(76). Normalized and smoothed coverage data used for ChIP-Seq plots was calculated using DANPOS2(77) and corrected using the fraction of deduplicated “spike-in” reads mapped to the mouse genome. Summary of ChIP-seq data is shown in Supplementary Table 2.

### Statistical analysis and plotting tools

All data was managed with Excel (Microsoft). All calculations and statistical tests not described in RNA-seq or ChIP-seq sections were run and plotted in Excel, except Mann-Whitney U tests run with an online calculator (http://www.statskingdom.com) and plots presented in Figures 2A and 5B created with BoxPlotR(78) and R respectively (www.R-project.org). For creation of the area proportional Venn diagrams presented in Figure 3B, the gene sets included in Supplementary Data 1 were first analyzed for overlaps by BioVenn(79), then plotted with Euler*APE*(80) and finally graphically adjusted (coloring, line formatting) in PowerPoint (Microsoft). Gene set enrichment analyses presented in Figure 3 were performed with PANTHER Overrepresentation Test(38, 39) (Supplementary Data 1). Gene set enrichment analyses presented in Figure 6C were performed with DAVID Functional Annotation Tool(81, 82) (Supplementary Data 3).

## Author Contributions

### Conceptualization

TB, PAZ, JS

### Experimental work

Preparation of mutant lines: TB, WS, AB, MA, PW, WD, MSz-L, Phenotypic analyses: TB, WD, MSz-L, RNA-seq/RT-qPCR: WS, TB, ChIP-seq/ChIP-qPCR: WS, TB, Protein-protein interactions: AB, TB, *In vitro* acetylation: WS, CL, AP-MS: NDW, TB

### Data analysis

TB, MK, PAZ

### Project administration

PAZ, JS

### Supervision

PAZ, JC (*in vitro* acetylation only), GDJ (AP-MS only)

### Writing and Editing

TB, PAZ with contributions from all authors

## Acknowledgements

This work was supported by the Polish National Science Center (NCN) grants 2016/21/B/NZ2/01757 and 2016/22/E/NZ2/00455 to PAZ, 2015/17/N/NZ1/00028 to W.S. and the Foundation for Polish Science grant (POIR.04.04.00-00-5C0F/17-00) to PAZ. The PhD fellowship of T.B. was covered by the International PhD Programme of the Foundation for Polish Science (MPD/2010/3) and KNOW RNA Research Centre in Poznan (01/KNOW2/2014). Research in JC lab was supported by a Foundation grant from the Canadian Institutes of Health Research (FDN-143314). JC holds a Canada Research Chair in Chromatin Biology and Molecular Epigenetics. The four-month stay of WS in JC group was funded by NCN grant 2017/24/T/NZ1/00165.

## Supplementary figure legends

**Supplementary Figure 1. DNA sequence analysis of the *Ateaf1-1* allele. A**, Gene structure diagram of the *AtEAF1* loci. Two different sequences targeted by gRNAs from the same genetic construct (#2) are marked with red and purple. Due to almost identical sequence of the two *AtEAF1* genes, each gRNA targeted both genes resulting in the total of four cut sites on each chromosome. **B**, A representative sequencing result of a genomic PCR clone from a single homozygous plant aligned with a fragment of the reference genomic sequences of *AtEAF1A and AtEAF1B.* All sequence alignment images in Supplementary Figures 1-3 were generated with Geneious 10 (Biomatters).

**Supplementary Figure 2. DNA sequence analysis of the *Atepl1a-2* allele. A**, Gene structure diagram of the *AtEPL1A* locus. Two different sequences targeted by gRNAs from the same genetic construct are marked with red and purple. **B**, A representative sequencing result of a genomic PCR clone from a single homozygous plant aligned with a fragment of the reference genomic sequence of *AtEPL1A.* The deletion starts in the exon II and ends in the intron IV (downstream from the second cut site), therefore, the protein product cannot be predicted based on the genomic sequence. The target sequences are framed and color-coded. Black arrow heads mark Cas9-gRNA cut sites. **C**, Sequences of 6 independent cDNA clones from a single homozygous mutant plant aligned with a fragment of the reference sequence of the *AtEPL1A* CDS. CDNA sequencing suggests that all transcripts are spliced in the same way which results in early translation termination. All sequence alignment images in Supplementary Figures 1-3 were generated with Geneious 10 (Biomatters).

**Supplementary Figure 3. DNA sequence analysis of the *Atepl1b-2* and *-3* alleles. A**, Gene structure diagram of the *AtEPL1B* locus. Two different sequences targeted by gRNAs from each of the two genetic constructs are marked with red and purple (*Atepl1b-2*) or orange and cyan (*Atepl1b-3*). **B**, A representative sequencing result of a genomic PCR clone from a single *Atepl1b-2* homozygous plant aligned with a fragment of the reference genomic sequence of *AtEPL1B.* The target sequences are framed and color-coded. Black arrow heads mark Cas9-gRNA cut sites. **C**, A representative sequencing result of a genomic PCR clone from a single *Atepl1b-3* homozygous plant aligned with a fragment of the reference genomic sequence of *AtEPL1B.* All sequence alignment images in Supplementary Figures 1-3 were generated with Geneious 10 (Biomatters).

**Supplementary Figure 4. The phenotype of *Atepl1-1* is more severe than the *Atyaf9a-1 Atyaf9b-2* mutant phenotype. A**, Quantitative description of 8-true leaf rosettes. Plants were grown on peat pellets (Jiffy) in the following conditions: 22°C, 16-hour light period, 40 μmol/(m^2^s) PAR, 70%. Error bars represent standard deviation. **B**, RT-qPCR analysis of relative gene expression in whole rosettes grown and collected as described above. Error bars represent standard deviation. Asterisks indicate significant differences relative to WT (two-tailed t-test, p < 0.05). **C**, Main stems of representative WT and *Atepl1-1* plants grown in the same conditions. **D**, Mean seed number per silique for siliques 11-20 collected from the main stem from three plants for each line. Error bars represent standard deviation.

**Supplementary Figure 5. Validation of the RNA-seq data by RT-qPCR.** Relative transcript levels of selected genes encoding proteins involved in chlorophyll biosynthesis were measured by RT-qPCR. Error bars represent standard deviation of the sample (3 biological replicates). The WT and *Atepl1-1* plants used for RNA extraction were grown in the same conditions as plants for RNA-seq (see Methods). If no significant difference between WT and *Atepl1-1* was detected by RNA-seq the data is not plotted (indicated by a black dot).

**Supplementary Figure 6. Early onset of the pathogen response-like transcriptomic pattern in *Atepl1-1*. A**, Representative images of plant material collected for the RT-qPCR analysis. Approximately 30 seedlings were used for each of the three biological replicates. **B**, RT-qPCR analysis of relative gene expression in 2-true leaf seedlings compared with RNA-seq data for 10-leaf rosettes obtained as described in Methods. Plants for RT-qPCR were grown on peat pellets (Jiffy) in the following conditions: 22°C, 16-hour light period, 40 μmol/(m^2^s) PAR, 70% RH. Error bars represent standard deviation.

**Supplementary Figure 7. ChIP-qPCR survey of chromatin perturbations in *Atepl1-2* and *Ateaf1-1* mutants**. Two *Atepl1*-hypoactive genes (*QQS*, *CHLI1*), one *Atepl1*-hyperactive stress-inducible gene (*PR1*) and one silent transposon (*Ta3*) were used to check for effects of *Atepl1* and *Ateaf1* mutations on chromatin. The graphs represent data for a single biological sample for each line. The error bars represent standard deviation of 3 technical replicates.

**Supplementary Figure 8. TSS ChIP-seq plots. A**, TSS-centered profiles of spike-in-normalized occupancy of H3, H3K9ac or H4K5ac averaged over all genes expressed in WT Col-0 (the same as in Fig. 5A), *Atepl1*-hypoactive or *Atepl1*-hyperactive genes. Vertical axes values are relative. Horizontal axes values represent chromosomal position relative to TSS (i.e. negative values represent promoter region whereas positive values represent gene body). **B**, As for A, but for H3K9ac. **C**, As for A, but for H4K5ac.

