## Supplementary Tables for "Nuclear encoded photosynthesis genes are specifically controlled by the NuA4 complex" (from another version of this preprint: version 1, 2019-11-18)

**Supplementary Table 1A.** Median chloroplast and cell size (corresponding to Figure 2A).

| **Median chloroplast projection surface area in µm² (n chloroplasts / replicate = 20)** | | | | | | | |
|---|---|---|---|---|---|---|---|
| Repl. # | WT | *Atepl1-1* | *Atepl1-2* | *Atepl1-3* | *Ateaf1-1* | *glk* | *Atepl1 glk* |
| 1 | 37.11 | 11.19 | 11.02 | 21.09 | 19.26 | 11.22 | 7.95 |
| 2 | 39.59 | 23.42 | 23.77 | 18.94 | 26.48 | 16.32 | 6.10 |
| 3 | 28.51 | 11.25 | 12.22 | 15.44 | 27.75 | 14.32 | 7.63 |
| 4 | 29.34 | 11.02 | 17.82 | 16.32 | 16.29 | 13.26 | 4.45 |
| 5 | 29.19 | 16.44 | 12.02 | 16.50 | 17.70 | 13.17 | 4.86 |
| 6 | 32.11 | 17.20 | 11.84 | 15.17 | 20.71 | 10.99 | 4.09 |
| 7 | 27.34 | 16.85 | 11.37 | 14.64 | 15.91 | 10.93 | 5.98 |
| 8 | 35.38 | 16.11 | 10.16 | 15.73 | 13.67 | 15.14 | 6.75 |
| **P values (two-tailed Mann Whitney U test)** | | | | | | | |
| | WT | *Atepl1-1* | *Atepl1-2* | *Atepl1-3* | *Ateaf1-1* | *glk* | *Atepl1 glk* |
| WT | | 0.000155 | 0.000155 | 0.000155 | 0.000311 | 0.000155 | 0.000155 |
| *Atepl1-1* | | | 0.645377 | 0.798446 | 0.130381 | 0.160528 | 0.000155 |
| *Atepl1-2* | | | | 0.064957 | 0.020668 | 0.878477 | 0.000155 |
| *Atepl1-3* | | | | | 0.234499 | 0.002953 | 0.000155 |
| *Ateaf1-1* | | | | | | 0.002953 | 0.000155 |
| *glk* | | | | | | | 0.000155 |
| *Atepl1 glk* | | | | | | | |

| **Median cell projection surface area in µm² (n cells / replicate = 9)** | | | | | | | |
|---|---|---|---|---|---|---|---|
| Repl. # | WT | *Atepl1-1* | *Atepl1-2* | *Atepl1-3* | *Ateaf1-1* | *glk* | *Atepl1 glk* |
| 1 | 3440.42 | 1357.28 | 1613.19 | 1610.72 | 1973.97 | 1796.00 | 979.53 |
| 2 | 3000.64 | 1856.62 | 1838.65 | 1248.41 | 2209.27 | 2395.61 | 612.16 |
| 3 | 1762.71 | 1168.82 | 1628.22 | 1271.33 | 2482.68 | 1904.75 | 759.08 |
| 4 | 2426.83 | 1157.92 | 1454.84 | 1423.20 | 1635.88 | 2228.77 | 2035.42 |
| 5 | 2658.77 | 1621.97 | 1040.92 | 1486.71 | 1791.99 | 1925.55 | 979.53 |
| 6 | 2528.16 | 1206.99 | 1370.12 | 1225.49 | 1661.38 | 1988.23 | 612.16 |
| 7 | 2226.82 | 1293.18 | 1558.29 | 1018.95 | 1539.85 | 2343.94 | 759.08 |
| 8 | 2284.62 | 1521.23 | 1299.72 | 1192.50 | 1691.55 | 2035.42 | 759.08 |
| **P values (two-tailed Mann Whitney U test)** | | | | | | | |
| | WT | *Atepl1-1* | *Atepl1-2* | *Atepl1-3* | *Ateaf1-1* | *glk* | *Atepl1 glk* |
| WT | | 0.000311 | 0.000311 | 0.000155 | 0.006993 | 0.049883 | 0.000311 |
| *Atepl1-1* | | | 0.441803 | 0.645377 | 0.004662 | 0.000311 | 0.010412 |
| *Atepl1-2* | | | | 0.104895 | 0.010412 | 0.000311 | 0.010412 |
| *Atepl1-3* | | | | | 0.000311 | 0.000155 | 0.010412 |
| *Ateaf1-1* | | | | | | 0.104895 | 0.004662 |
| *glk* | | | | | | | 0.001865 |
| *Atepl1 glk* | | | | | | | |

**Supplementary Table 1B.** Total chlorophyll and chlorophyll a / b ratio (corresponding to Figure 2B).

| **Total Chlorophyll (nmol total chlorophyll / mg fresh weight)** | | | | | | | |
|---|---|---|---|---|---|---|---|
| | WT | *Atepl1-1* | *Atepl1-2* | *Atepl1-3* | *Ateaf1-1* | *glk* | *Atepl1 glk* |
| | 1.36 | 0.58 | 0.55 | 0.60 | 0.62 | 0.45 | 0.30 |
| | 1.29 | 0.52 | 0.54 | 0.61 | 0.60 | 0.39 | 0.33 |
| | 1.51 | 0.58 | 0.48 | 0.60 | 0.61 | 0.52 | 0.31 |
| Mean | 1.39 | 0.56 | 0.53 | 0.61 | 0.61 | 0.45 | 0.31 |
| alpha | 0.05 | 0.05 | 0.05 | 0.05 | 0.05 | 0.05 | 0.05 |
| SD | 0.11 | 0.03 | 0.04 | 0.01 | 0.01 | 0.07 | 0.02 |
| n | 3 | 3 | 3 | 3 | 3 | 3 | 3 |
| CI 95% | 0.28 | 0.08 | 0.10 | 0.02 | 0.03 | 0.16 | 0.04 |
| **P values (two-tailed T-test)** | | | | | | | |
| | WT | *Atepl1-1* | *Atepl1-2* | *Atepl1-3* | *Ateaf1-1* | *glk* | *Atepl1 glk* |
| WT | | 0.00026 | 0.00023 | 0.00028 | 0.00029 | 0.00024 | 0.00008 |
| *Atepl1-1* | | | 0.29529 | 0.09824 | 0.07235 | 0.06581 | 0.00030 |
| *Atepl1-2* | | | | 0.02606 | 0.02122 | 0.18212 | 0.00086 |
| *Atepl1-3* | | | | | 0.41410 | 0.01673 | 0.00001 |
| *Ateaf1-1* | | | | | | 0.01478 | 0.00001 |
| *glk* | | | | | | | 0.01964 |
| *Atepl1 glk* | | | | | | | |
| **Chlorophyll a / b ratio** | | | | | | | |
| | WT | *Atepl1-1* | *Atepl1-2* | *Atepl1-3* | *Ateaf1-1* | *glk* | *Atepl1 glk* |
| | 3.34 | 3.21 | 2.91 | 3.44 | 3.22 | 4.92 | 3.67 |
| | 3.31 | 3.12 | 3.27 | 2.98 | 3.22 | 4.66 | 3.76 |
| | 3.26 | 3.07 | 2.96 | 3.01 | 3.32 | 4.85 | 3.49 |
| Mean | 3.30 | 3.13 | 3.05 | 3.14 | 3.25 | 4.81 | 3.64 |
| alpha | 0.05 | 0.05 | 0.05 | 0.05 | 0.05 | 0.05 | 0.05 |
| SD | 0.04 | 0.07 | 0.19 | 0.26 | 0.06 | 0.14 | 0.14 |
| n | 3 | 3 | 3 | 3 | 3 | 3 | 3 |
| CI 95% | 0.10 | 0.19 | 0.48 | 0.64 | 0.14 | 0.34 | 0.35 |
| **P values (two-tailed T-test)** | | | | | | | |
| | WT | *Atepl1-1* | *Atepl1-2* | *Atepl1-3* | *Ateaf1-1* | *glk* | *Atepl1 glk* |
| WT | | 0.02508 | 0.08391 | 0.34148 | 0.27991 | 0.00005 | 0.01634 |
| *Atepl1-1* | | | 0.49917 | 0.95560 | 0.09151 | 0.00005 | 0.00525 |
| *Atepl1-2* | | | | 0.62631 | 0.14550 | 0.00020 | 0.01239 |
| *Atepl1-3* | | | | | 0.50408 | 0.00057 | 0.04253 |
| *Ateaf1-1* | | | | | | 0.00005 | 0.01160 |
| *glk* | | | | | | | 0.00048 |
| *Atepl1 glk* | | | | | | | |

**Supplementary Table 1C.** Rosette size in one-week intervals (corresponding to Figure 2C).

| **Log$_{10}$ of rosette radius in mm** | | | | | | | |
|---|---|---|---|---|---|---|---|
| **Week 2** | | | | | | | |
| Plant # | WT | *Atepl1-1* | *Atepl1-2* | *Atepl1-3* | *Ateaf1-1* | *glk* | *Atepl1 glk* |
| 1 | 0.89 | 0.18 | 0.20 | 0.38 | 0.44 | 0.77 | 0.36 |
| 2 | 0.87 | 0.10 | 0.16 | 0.31 | 0.47 | 0.73 | 0.20 |
| 3 | 0.85 | 0.10 | 0.35 | 0.21 | 0.39 | 0.80 | 0.33 |
| 4 | 1.01 | 0.41 | 0.37 | -0.05 | 0.19 | 0.72 | 0.34 |
| 5 | 0.78 | 0.36 | 0.33 | 0.35 | 0.42 | 0.74 | 0.14 |
| 6 | 0.64 | 0.28 | 0.11 | 0.31 | 0.40 | 0.72 | 0.25 |
| 7 | 0.78 | 0.39 | 0.36 | 0.28 | 0.40 | 0.73 | 0.12 |
| 8 | 0.76 | 0.29 | 0.25 | 0.40 | 0.29 | 0.68 | 0.02 |
| 9 | 0.83 | 0.38 | 0.19 | 0.20 | 0.32 | 0.70 | 0.23 |
| 10 | 0.87 | 0.13 | 0.15 | 0.28 | 0.33 | 0.57 | 0.16 |
| 11 | | | 0.20 | | | | |
| 12 | | | 0.17 | | | | |
| 13 | | | 0.33 | | | | |
| 14 | | | 0.33 | | | | |
| 15 | | | 0.21 | | | | |
| 16 | | | 0.27 | | | | |
| 17 | | | 0.36 | | | | |
| 18 | | | 0.37 | | | | |
| 19 | | | 0.34 | | | | |
| 20 | | | 0.22 | | | | |
| Mean | 0.83 | 0.26 | 0.26 | 0.27 | 0.37 | 0.72 | 0.22 |
| alpha | 0.05 | 0.05 | 0.05 | 0.05 | 0.05 | 0.05 | 0.05 |
| SD | 0.10 | 0.13 | 0.09 | 0.13 | 0.08 | 0.06 | 0.11 |
| n | 10 | 10 | 20 | 10 | 10 | 10 | 10 |
| CI 95% | 0.07 | 0.09 | 0.04 | 0.09 | 0.06 | 0.04 | 0.08 |
| **P values (two-tailed T-test)** | | | | | | | |
| | WT | *Atepl1-1* | *Atepl1-2* | *Atepl1-3* | *Ateaf1-1* | *glk* | *Atepl1 glk* |
| WT | | 1.32E-09 | 9.88E-16 | 2.29E-09 | 1.19E-09 | 7.05E-03 | 1.13E-10 |
| *Atepl1-1* | | | 9.55E-01 | 9.12E-01 | 4.21E-02 | 5.48E-09 | 3.98E-01 |
| *Atepl1-2* | | | | 9.16E-01 | 4.64E-03 | 1.00E-14 | 2.03E-01 |
| *Atepl1-3* | | | | | 6.06E-02 | 1.08E-08 | 3.47E-01 |
| *Ateaf1-1* | | | | | | 3.31E-09 | 3.05E-03 |
| *glk* | | | | | | | 2.50E-10 |
| *Atepl1 glk* | | | | | | | |

| **Log$_{10}$ of rosette radius in mm** | | | | | | | |
|---|---|---|---|---|---|---|---|
| **Week 3** | | | | | | | |
| Plant # | WT | *Atepl1-1* | *Atepl1-2* | *Atepl1-3* | *Ateaf1-1* | *glk* | *Atepl1 glk* |
| 1 | 1.35 | 0.51 | 0.47 | 0.58 | 0.67 | 1.16 | 0.49 |
| 2 | 1.45 | 0.24 | 0.36 | 0.65 | 0.85 | 1.11 | 0.29 |
| 3 | 1.38 | 0.51 | 0.71 | 0.49 | 0.82 | 1.14 | 0.46 |
| 4 | 1.51 | 0.71 | 0.60 | 0.24 | 0.54 | 1.15 | 0.47 |
| 5 | 1.40 | 0.58 | 0.48 | 0.59 | 0.78 | 1.11 | 0.32 |
| 6 | 1.30 | 0.46 | 0.19 | 0.53 | 0.79 | 1.10 | 0.42 |
| 7 | 1.38 | 0.72 | 0.60 | 0.53 | 0.71 | 1.10 | 0.28 |
| 8 | 1.34 | 0.55 | 0.43 | 0.65 | 0.63 | 1.06 | 0.18 |
| 9 | 1.39 | 0.69 | 0.42 | 0.48 | 0.63 | 1.10 | 0.42 |
| 10 | 1.38 | 0.31 | 0.38 | 0.59 | 0.61 | 0.98 | 0.39 |
| 11 | | | 0.48 | | | | |
| 12 | | | 0.42 | | | | |
| 13 | | | 0.60 | | | | |
| 14 | | | 0.69 | | | | |
| 15 | | | 0.52 | | | | |
| 16 | | | 0.49 | | | | |
| 17 | | | 0.64 | | | | |
| 18 | | | 0.53 | | | | |
| 19 | | | 0.57 | | | | |
| 20 | | | 0.43 | | | | |
| Mean | 1.39 | 0.53 | 0.50 | 0.53 | 0.70 | 1.10 | 0.37 |
| alpha | 0.05 | 0.05 | 0.05 | 0.05 | 0.05 | 0.05 | 0.05 |
| SD | 0.06 | 0.16 | 0.12 | 0.12 | 0.10 | 0.05 | 0.10 |
| n | 10 | 10 | 20 | 10 | 10 | 10 | 10 |
| CI 95% | 0.04 | 0.11 | 0.06 | 0.09 | 0.07 | 0.04 | 0.07 |
| **P values (two-tailed T-test)** | | | | | | | |
| | WT | *Atepl1-1* | *Atepl1-2* | *Atepl1-3* | *Ateaf1-1* | *glk* | *Atepl1 glk* |
| WT | | 4.46E-12 | 6.08E-19 | 6.20E-14 | 3.48E-13 | 6.89E-10 | 2.54E-16 |
| *Atepl1-1* | | | 5.76E-01 | 9.64E-01 | 9.95E-03 | 2.91E-09 | 1.66E-02 |
| *Atepl1-2* | | | | 4.93E-01 | 1.16E-04 | 1.12E-14 | 8.54E-03 |
| *Atepl1-3* | | | | | 2.99E-03 | 4.79E-11 | 4.26E-03 |
| *Ateaf1-1* | | | | | | 1.96E-09 | 8.22E-07 |
| *glk* | | | | | | | 5.80E-14 |
| *Atepl1 glk* | | | | | | | |

| Log10 of rosette radius in mm | | | | | | | |
|---|---|---|---|---|---|---|---|
| **Week 4** | | | | | | | |
| Plant # | WT | *Atepl1-1* | *Atepl1-2* | *Atepl1-3* | *Ateaf1-1* | *glk* | *Atepl1 glk* |
| 1 | 1.69 | 0.87 | 0.82 | 0.88 | 1.00 | 1.48 | 0.73 |
| 2 | 1.75 | 0.69 | 0.52 | 0.86 | 1.06 | 1.39 | 0.56 |
| 3 | 1.71 | 0.82 | 0.82 | 0.72 | 1.06 | 1.49 | 0.69 |
| 4 | 1.74 | 0.96 | 0.82 | 0.65 | 0.88 | 1.49 | 0.69 |
| 5 | 1.70 | 0.88 | 0.83 | 0.86 | 1.02 | 1.45 | 0.68 |
| 6 | 1.69 | 0.89 | 0.28 | 0.82 | 1.04 | 1.42 | 0.61 |
| 7 | 1.71 | 0.94 | 0.83 | 0.79 | 0.92 | 1.44 | 0.54 |
| 8 | 1.69 | 0.82 | 0.73 | 0.93 | 0.90 | 1.42 | 0.28 |
| 9 | 1.67 | 0.94 | 0.76 | 0.85 | 0.93 | 1.42 | 0.60 |
| 10 | 1.68 | 0.62 | 0.77 | 0.89 | 0.84 | 1.35 | 0.69 |
| 11 | | | 0.86 | | | | |
| 12 | | | 0.69 | | | | |
| 13 | | | 0.87 | | | | |
| 14 | | | 0.87 | | | | |
| 15 | | | 0.80 | | | | |
| 16 | | | 0.84 | | | | |
| 17 | | | 0.86 | | | | |
| 18 | | | 0.83 | | | | |
| 19 | | | 0.93 | | | | |
| 20 | | | 0.70 | | | | |
| Mean | 1.70 | 0.84 | 0.77 | 0.83 | 0.96 | 1.43 | 0.61 |
| alpha | 0.05 | 0.05 | 0.05 | 0.05 | 0.05 | 0.05 | 0.05 |
| SD | 0.03 | 0.11 | 0.15 | 0.09 | 0.08 | 0.04 | 0.13 |
| n | 10 | 10 | 20 | 10 | 10 | 10 | 10 |
| CI 95% | 0.02 | 0.08 | 0.07 | 0.06 | 0.06 | 0.03 | 0.09 |
| | WT | *Atepl1-1* | *Atepl1-2* | *Atepl1-3* | *Ateaf1-1* | *glk* | *Atepl1 glk* |
| WT | | 5.33E-15 | 4.80E-18 | 4.32E-17 | 3.77E-16 | 3.23E-12 | 9.90E-16 |
| *Atepl1-1* | | | 1.82E-01 | 6.78E-01 | 1.26E-02 | 7.56E-12 | 4.01E-04 |
| *Atepl1-2* | | | | 2.97E-01 | 5.85E-04 | 4.03E-14 | 5.73E-03 |
| *Atepl1-3* | | | | | 1.44E-03 | 9.84E-14 | 3.49E-04 |
| *Ateaf1-1* | | | | | | 4.03E-12 | 8.08E-07 |
| *glk* | | | | | | | 2.47E-13 |
| *Atepl1 glk* | | | | | | | |

**Supplementary Table 2.** ChIP-seq summary.

| Sample | Antibody | Total reads | Mapped reads | Duplicates | Analyzed reads | Mouse reads | Fraction of mouse | Spike-in |
|---|---|---|---|---|---|---|---|---|
| epl2_H3 | H3 | 21368578 | 19748913 | 5495889 | 14253024 | 537809 | 0.0364 | 0.0324 |
| epl3_H3 | H3 | 19300672 | 17869582 | 5371442 | 12498140 | 399292 | 0.0310 | |
| epl4_H3 | H3 | 20967261 | 18446233 | 6138110 | 12308123 | 380419 | 0.0300 | |
| WT2_1_H3 | H3 | 22194295 | 21392573 | 5947892 | 15444681 | 522769 | 0.0327 | 0.0373 |
| WT3_H3 | H3 | 20235352 | 19536076 | 6273084 | 13262992 | 502727 | 0.0365 | |
| WT4_H3 | H3 | 22392052 | 21504402 | 6620439 | 14883963 | 660968 | 0.0425 | |
| epl2_H3K9ac | H3K9ac | 6482922 | 5300353 | 1456456 | 3843897 | 127218 | 0.0320 | Not used |
| epl3_H3K9ac | H3K9ac | 17265676 | 13782719 | 4575676 | 9207043 | 145810 | 0.0156 | 0.0172 |
| epl4_H3K9ac | H3K9ac | 19219574 | 13684182 | 3461175 | 10223007 | 195414 | 0.0188 | |
| WT2_1_H3K9ac | H3K9ac | 18074380 | 16849044 | 4838713 | 12010331 | 219056 | 0.0179 | 0.0185 |
| WT3_H3K9ac | H3K9ac | 18639713 | 16844507 | 5677908 | 11166599 | 171393 | 0.0151 | |
| WT4_H3K9ac | H3K9ac | 18395451 | 16431382 | 5497196 | 10934186 | 251640 | 0.0225 | |
| epl2_H4K5ac | H4K5ac | 19861888 | 10439005 | 3263945 | 7175060 | 1089966 | 0.1319 | 0.1338 |
| epl3_H4K5ac | H4K5ac | 19481167 | 10547446 | 2839972 | 7707474 | 787961 | 0.0928 | |
| epl4_H4K5ac | H4K5ac | 21824904 | 8028253 | 2410606 | 5617647 | 1206501 | 0.1768 | |
| WT2_1_H4K5ac | H4K5ac | 18713901 | 16778202 | 4544722 | 12233480 | 454494 | 0.0358 | 0.0634 |
| WT2_H4K5ac | H4K5ac | 17315277 | 15632424 | 4949223 | 10683201 | 643418 | 0.0568 | |
| WT4_H4K5ac | H4K5ac | 17765646 | 15169934 | 4889503 | 10280431 | 1110910 | 0.0975 | |

**Supplementary Table 3.** Oligonucleotides used in this work.

| Template | Name | Sequence | Source | Notes |
|---|---|---|---|---|
| **Genotyping** | | | | |
| *Ateaf1-1* | F | GGGGGATCCTCAGCCTCT | This work | |
| *Ateaf1-1* | R | GGCCTTTTATTTTTCTGCCCCT | This work | |
| *atepl1a-1* | LP | GCCATGAAATCCCTCTTTTTC | http://signal.salk.edu/tdnaprimers.2.html | |
| *atepl1a-1* | RP | TACTCCTACCCTTGGTTTGCC | http://signal.salk.edu/tdnaprimers.2.html | |
| *atepl1b-1* | LP | AGCGTAAAAGATGGCAGAAGC | http://signal.salk.edu/tdnaprimers.2.html | |
| *atepl1b-1* | RP | TACCAGTCTGTGAAGCCATCC | http://signal.salk.edu/tdnaprimers.2.html | |
| *atepl1a-2* | F | CCCGATTTTTGATTAGCCGGTAC | This work | |
| *atepl1a-2* | R | CCATCTACCACACAACAAAGCTC | This work | |
| *atepl1b-2* | F | TTCGTCCTGTCTTATTGGAGCAA | This work | |
| *atepl1b-2* | R | CCAATCCGCCTCAACGTTTAAAA | This work | |
| *atepl1b-3* | F | AGCTCTTGGAAGATAGCTTGGC | This work | |
| *atepl1b-3* | R | ATTGCATACACCATTGGTGATGG | This work | |
| T-DNA | LBb1.3 | ATTTTGCCGATTTCGGAAC | http://signal.salk.edu/tdnaprimers.2.html | |
| T-DNA | LB3 | TAGCATCTGAATTTCATAACCAATCTCGATACAC | http://signal.salk.edu/tdnaprimers.2.html | |
| *atglk1.1* | Spm5 | CGGGATCCGACACTCTTTAATTAACTGACACTC | Fitter et al., 2002 | |
| *atglk1.1* | glk1f | GGGTATTCGGGTCGGGTATT | This work | |
| *atglk1.1* | glk1r | CCCTTCGTTGTTACTGATCCG | This work | |
| *atglk2.1* | Spm1 | CCTATTTCAGTAAGAGTGTGGGGTTTTGG | Fitter et al., 2002 | |
| *atglk2.1* | GLK2F | CTCTCCTATCTCTCAACCACA | Fitter et al., 2002 | |
| *atglk2.1* | GLK2R | AGATGTTTCCGATGTGACCT | Fitter et al., 2002 | |
| **gRNA mutagenesis** | | | | |
| gRNA vector U3 | Ateaf1 pair 1 g1 | CCATTATCAGTGAGTGCATTGACCCTTGATGCTTTCTATGCA | This work | |
| gRNA vector U6 | Ateaf1 pair 1 g2 | GAGGAATTCCAGAGCACAACAATCACTACTTCGACTCTAGCTGT | This work | |
| gRNA vector U6 | Ateaf1 pair 2 g1 | ATTGCTTGGTGGTCTTTTGCAATCACTACTTCGACTCTAGCTGT | This work | |
| gRNA vector U3 | Ateaf1 pair 2 g2 | ATCACGACCTCCAATAAATTGACCCTTGATGCTTTCTATGCA | This work | |
| gRNA vector U6 | Ateaf1 pair 3 g1 | CCAGGAACCATGCGATGTCCAATCACTACTTCGACTCTAGCTGT | This work | |
| gRNA vector U3 | Ateaf1 pair 3 g2 | TGACATGGGTGTGTTCCTGTGACCCTTGATGCTTTCTATGCA | This work | |
| gRNA vector U3 | Atepl1b-1 g1 | GTGGACGGGGTCGAAAAGATGACCCTTGATGCTTTCTATGCA | This work | |

| | | | | |
|---|---|---|---|---|
| gRNA vector U6 | Atepl1b-1 g2 | AGCTCAGCCCGAGCTGCAAACAAT CACTACTTCGACTCTAGCTGT | This work | |
| gRNA vector U6 | Atepl1b-2 g1 | GCCACGGCCATTAGACATTCAATC ACTACTTCGACTCTAGCTGT | This work | |
| gRNA vector U6 | Atepl1b-2 g2 | AGCTCAGACCGAGCTGCATACAAT CACTACTTCGACTCTAGCTGT | This work | |
| gRNA vector U3 | Atepl1b-3 g1 | GGTCAACCTCCAAGTCGGTTGACC CTTGATGCTTTCTATGCA | This work | |
| gRNA vector U3 | Atepl1b-3 g2 | AATCACCGGTCTACCAACTTGACC CTTGATGCTTTCTATGCA | This work | |
| **RT-PCR / RT-qPCR / ChIP-qPCR** | | | | |
| AT3G01500 | CA1_F | TGTGTCCATCACACGTTCTGG | (Woodson et al., 2013) | |
| AT3G01500 | CA1_R | GGACCACGAAGGCATCTCCT | (Woodson et al., 2013) | |
| AT1G16690 | AtEPL1A_F | TTGTTGACAATGAGGTGCAGC | This work | Primer # 1 in Fig. 1 B |
| AT1G16690 | AtEPL1A_R1 | TACTGAGGTGTTGGGATTTCCG | This work | Primer # 2 in Fig. 1 B |
| AT1G16690 | AtEPL1A_R2 | AGGCCCATGAGGCCATGGTGCTAT GTAGAGAGTGTT | This work | Primer # 3 in Fig. 1 B |
| AT1G79020 | AtEPL1B_F | TGACAATGAGGTGGCTCCAG | This work | Primer # 4 in Fig. 1 B |
| AT1G79020 | AtEPL1B_R1 | AAGCAGGCTGACCAAAAGTC | This work | Primer # 5 in Fig. 1 B |
| AT1G79020 | AtEPL1B_R2 | TTTGATGCCTCCGGGATGAG | This work | Primer # 6 in Fig. 1 B |
| AT1G58290 | HEMA1_F | ACTCTGCAGCTGATCGATATACA | This work | |
| AT1G58290 | HEMA1_R | AGCAAGCTTCTCACGCATCT | This work | |
| AT1G29910, AT1G29920, AT1G29930 | LHCB1_F | AGCTCAAGAACGGAAGATTGG | (Kakizaki et al., 2009) | |
| AT1G29910, AT1G29920, AT1G29930 | LHCB1_R | GCCAAATGGTCAGCAAGGTT | (Kakizaki et al., 2009) | |
| AT2G05100, AT2G05070, AT3G27690 | LHCB2_F | CGTGTGGTTCAAGGCAGG | This work | |
| AT2G05100, AT2G05070, AT3G27690 | LHCB2_R | GATCAAGTTAGGGTTTCCGAGG | This work | |
| AT5G54270 | LHCB3_F | TCCTTGGCCAGACTAAAGCC | This work | |
| AT5G54270 | LHCB3_R | TACCAAAGATCATTTCCCATAGTG T | This work | |
| AT1G67090 | RBCS1A_F | AATTTCCGGACTTAACGTTTGTTT | (Izumi et al., 2012) | |
| AT1G67090 | RBCS1A_R | CATCAGACAGTTGAGAATCCGATA GA | (Izumi et al., 2012) | |
| AT5G38410 | RBCS3B_F | CCTATTGTCTGTGTTCTTTTTCTC TTTATG | (Izumi et al., 2012) | |
| AT5G38410 | RBCS3B_R | TCAAGACGCACGGATATATAAATT ACA | (Izumi et al., 2012) | |
| AT4G05320 | UBQ10_F | GGCCTTGTATAATCCCTGATGAAT AAG | This work | |
| AT4G05320 | UBQ10_R | AAAGAGATAACAGGAACGGAAACA TAG | This work | |
| AT3G57260 | BGL2qF | TCGCTGGATGTTTTGTTGTTTGT | This work | |
| AT3G57260 | BGL2qR | TATGCTACGGGATGCTAGGC | This work | |

| | | | |
|---|---|---|---|
| AT3G48090 | EDS1qF | AGCGCTCGGTTAGGTAAGC | This work |
| AT3G48090 | EDS1qR | GCGTTTGAAGCTCTTACCGG | This work |
| AT4G19660 | NPR4qF | GTGAAACTTATAGATGAAGATGGC TCT | This work |
| AT4G19660 | NPR4qR | AAACAGAGTCTTTGAGTGATACTT CTG | This work |
| AT3G52430 | PAD4qF | GCCGGGATTACATACGTTGC | This work |
| AT3G52430 | PAD4qR | AGCCGGGGAAAAGAACATCT | This work |
| AT5G44420 | PDF1qF | GAGCAGCGAAGATAAGGGTGA | This work |
| AT5G44420 | PDF1qR | TCACACAACACATACATCTATACA TTGA | This work |
| AT2G14610 | PR1qF | TTGTGAACCCTTAGATAATCTTGT GG | This work |
| AT2G14610 | PR1qR | TTTAATCGTCTTTGTAGCTCTTGT AGG | This work |
| AT1G75040 | PR5qF | GCATTGCTGTTATGGCCACA | This work |
| AT1G75040 | PR5qR | TCAAATCCTCCATCGCCGAG | This work |
| AT5G14930 | SAG101qF | AAGCTTGTGGTGACCTCTGG | This work |
| AT5G14930 | SAG101qR | TGTAGACCTGGATCATGATTCGAA | This work |
| AT1G58290 | HEMA1qF | ACAGTAGAACGCTGAGCGAG | This work |
| AT1G58290 | HEMA1qR | GTTGTTCCGCCATTGCCTTA | This work |
| AT5G63570 | GSA1qF | GCGACACAGAGAAGTTTGGT | This work |
| AT5G63570 | GSA1qR | CCTACTCAGTACCCTCTCAGC | This work |
| AT1G69740 | HEMB1qF | CGATGATCAAAGCAGGGGGA | This work |
| AT1G69740 | HEMB1q | GAGGATGATGTCCGCTCCAG | This work |
| AT5G08280 | HEMCqF | AAGATGGGAAAAGACGCGGG | This work |
| AT5G08280 | HEMCqR | CCGTGTCACATCTCTTCCTCC | This work |
| AT2G26540 | HEMDqF | GTGGGTCGAGAGCATAATGGA | This work |
| AT2G26540 | HEMDqR | GGCACTGACCAAACTCAAAGT | This work |
| AT2G40490 | HEME2qF | ACTGTGAAGAAAGCTGGGAGA | This work |
| AT2G40490 | HEME2qR | GCAACCTCAAAGAAGTGTGCTA | This work |
| AT1G03475 | HEMF1qF | GAAGAGGGGACCGAAGAGTG | This work |
| AT1G03475 | HEMF1qR | TCACTCTTCTCACAACAATGGGA | This work |
| AT4G01690 | HEMG1qF | CGGGCTACGAAGGGCTATTT | This work |
| AT4G01690 | HEMG1qR | TGACCTCAATCGCGGTTTCA | This work |
| AT1G08520 | CHLDqF | TGTAATCTCGGCCACCACTAG | This work |
| AT1G08520 | CHLDqR | TGTTTAGTGACTGATGTGTTGCA | This work |
| AT5G13630 | CHLHqF | AATGGTCGTGGCTACTGGG | This work |
| AT5G13630 | CHLHqR | GGCTTATATCCCATTGTTTATCGA TCG | This work |
| AT3G59400 | GUN4qF | GGATTTCTCTTCAATGTCAGTTCC A | This work |
| AT3G59400 | GUN4qR | TGATCAACCAAGAAAGAGTCATGT | This work |
| AT4G18480 | CHLI1qF | ACTAACAGAGCAGCAAAAGCAC | This work |
| AT4G18480 | CHLI1qR | TCCTCAGACGGTGCCTTAAG | This work |
| AT4G25080 | CHLMqF | CCAGGGCCATCAAAAGCTAC | This work |
| AT4G25080 | CHLMqR | ACTGTGTGGTAGTGAGTCCTC | This work |
| AT3G56940 | CRD1qF | GCTATGAGAAGCTGCTGGCT | This work |
| AT3G56940 | CRD1qR | GGCTCAAACTCGGCGAAATC | This work |
| AT5G54190 | PORAqF | CTTGACGAAGTCGGGAGTGT | This work |

| | | | |
|---|---|---|---|
| AT5G54190 | PORAqR | GAGCTTCTCGCTGACTTCCC | This work |
| AT4G27440 | PORBqF | CAAGCTTGACGAAATCAGGGG | This work |
| AT4G27440 | PORBqR | TCCCACACTTTACGAGCCTTC | This work |
| AT1G03630 | PORCqF | GTCTTGGGAAGTCAGGAGTGT | This work |
| AT1G03630 | PORCqR | TTCTCGCTAACCTCCCACAG | This work |
| AT5G18660 | DVRqF | ACTGGAGAGTATAGCGAGGAGA | This work |
| AT5G18660 | DVRqR | GTTCACCGAGTTCTTGACCC | This work |
| AT1G44446 | CAOqF | ACGGATGTTAAACGGAGCAAAC | This work |
| AT1G44446 | CAOqR | ACAGTTAGCCGGAGAAAGGT | This work |
